# FlyPredictome: A structural atlas of predicted protein-protein interactions in *Drosophila*

**DOI:** 10.64898/2026.04.14.718529

**Authors:** Ah-Ram Kim, Aram Comjean, Austin Veal, Jonathan Rodiger, Myeonghoon Han, Yanhui Hu, Norbert Perrimon

## Abstract

Protein-protein interactions (PPIs) are fundamental to cellular function. Yet most *Drosophila* PPIs remain structurally uncharacterized despite the wealth of genetic and biochemical data available for this organism. Here we present FlyPredictome, a structural interactome based on 1.5 million pairwise AlphaFold-Multimer predictions. Using a local confidence metric that isolates the binding site from non-interacting regions, we systematically assess experimentally reported *Drosophila* PPIs and predict direct binding interfaces at residue-level resolution. Testing their functional relevance, we find that phenotype-associated missense mutations are enriched at predicted interaction interfaces. Building on these predictions, we construct an evidence-supported PPI network, revealing modular organization from signaling pathways to individual protein complexes. We further predict nearly 400 higher-order assemblies drawn from these modules with AlphaFold3, many matching cryo-EM structures and some extending beyond them. FlyPredictome is available as an open, interactive database that maps interactions to residue-level binding surfaces, providing confidence-ranked candidates for interaction discovery in *Drosophila*.

## INTRODUCTION

*Drosophila melanogaster* has been a model for dissecting fundamental biological mechanisms, including cell signaling and development. Forward and reverse genetic screens have mapped the core components and epistatic hierarchies of conserved pathways, including Notch, Wnt/Wingless, Hedgehog, JAK-STAT, Hippo, TGF-β, Toll, and receptor tyrosine kinase cascades, establishing a rich catalog of functional relationships among signaling proteins (reviewed in St Johnston, 2002; Bier, 2005; Perrimon et al., 2012; Attrill et al., 2024). Yet genetic interaction does not equate to physical interaction: a suppressor or enhancer relationship establishes functional linkage but does not reveal whether two proteins bind directly, which domains mediate the contact, or which residues form the interface. Even for most genetically defined pathway connections, the structural basis of the underlying physical interaction remains largely unknown.

Large-scale experimental approaches have expanded the catalog of physical interactions. The FlyBi yeast two-hybrid (Y2H) screen tested >10,000 *Drosophila* proteins in all-by-all screens, identifying ∼8,700 binary interactions (Tang et al., 2023), while affinity purification-mass spectrometry (AP-MS) studies (DPiM, DPiM2) mapped co-complex associations across thousands of baits (Guruharsha et al., 2011; Bhat et al., 2024). However, these datasets remain incomplete in coverage and provide no structural information about the interaction interface.

Recent advances in deep learning-based structure prediction, including AlphaFold (Jumper et al., 2021) and RoseTTAFold (Baek et al., 2021), have enabled prediction of three-dimensional structures of protein complexes. These methods have been applied at proteome scale to generate structural interactomes across multiple organisms (Humphreys et al., 2021; Burke et al., 2023; Schmid and Walter, 2025; Schmid et al., 2025; Zhang et al., 2025; Han et al., 2026; Qi et al., 2026). Although structural predictions have been reported for selected *Drosophila* proteins and protein pairs (Kawaguchi et al., 2025; Peng and Zhao, 2026) and for STRING-nominated pairs in the AlphaFold Database (Han et al., 2026), structural coverage of experimentally reported *Drosophila* interactions remains incomplete.

Here, we present FlyPredictome, a structural atlas of the *Drosophila melanogaster* protein interactome based on 1.5 million pairwise AlphaFold-Multimer predictions. We score predictions using a local confidence metric (integrated Local Interaction Score, iLIS), evaluate recovery of experimentally reported *Drosophila* protein-protein interactions (PPIs), and provide structural models at residue-level resolution. We further test predicted interfaces by demonstrating that phenotype-associated missense mutations are enriched at predicted interaction sites. We organize evidence-supported predictions into a hierarchically clustered PPI network of over 21,000 interactions, revealing functional organization from pathway-level modules down to individual protein complexes. We also show that assemblies drawn from these modules fold into higher-order complexes with AlphaFold3, many matching available cryo-EM structures, including those solved in non-fly orthologs, and some extending to assemblies not yet experimentally resolved. Finally, we provide the FlyPredictome database and an interactive network viewer, in which each protein links to the residues it uses to bind its partners, as open resources for the community.

## RESULTS

### A local confidence metric for protein interactions

To evaluate the ability of AlphaFold-Multimer (AFM v2.3; Evans et al., 2021; Google DeepMind, 2022; Methods) to predict PPIs, we tested it on the Positive Reference Sets (PRS) from proteome-scale yeast (Yu et al., 2008), fly (Tang et al., 2023), and human (Braun et al., 2009) Y2H screens. Several well-established PPIs, including p53-MDM2 (human), Emc-Da (fly), and SEC9-SSO1 (yeast), received low scores using the interface predicted TM-score (ipTM), a widely used metric for evaluating AFM prediction confidence, despite forming clearly defined local interfaces visible in the Predicted Aligned Error (PAE) heatmaps (**Figure S1A**).

We previously introduced the Local Interaction Score (LIS) framework (Kim et al., 2024), which scores the confident interaction domain (residue pairs with inter-chain PAE ≤ 12 Å, rescaled so that lower PAE yields a higher score) to evaluate PPIs. However, in some cases, the inter-chain residue pairs that AFM positions most confidently (low PAE) are not in direct contact, often terminal segments placed far from the partner, so a high LIS can arise without a genuine contact interface (**Figure S1B**). To suppress such false positives, we developed iLIS, the geometric mean of LIS and cLIS (contact LIS), the latter restricting scoring to residues in direct intermolecular contact (Cβ-Cβ ≤ 8 Å within the PAE ≤ 12 Å domain); because this geometric mean collapses toward zero when contacts are absent (cLIS = 0), it penalizes these cases and integrates domain- and contact-level confidence into a single, balanced score (**Figure 1A**; alternative formulations perform similarly; **Figure S2** and Methods).

**Figure 1.**
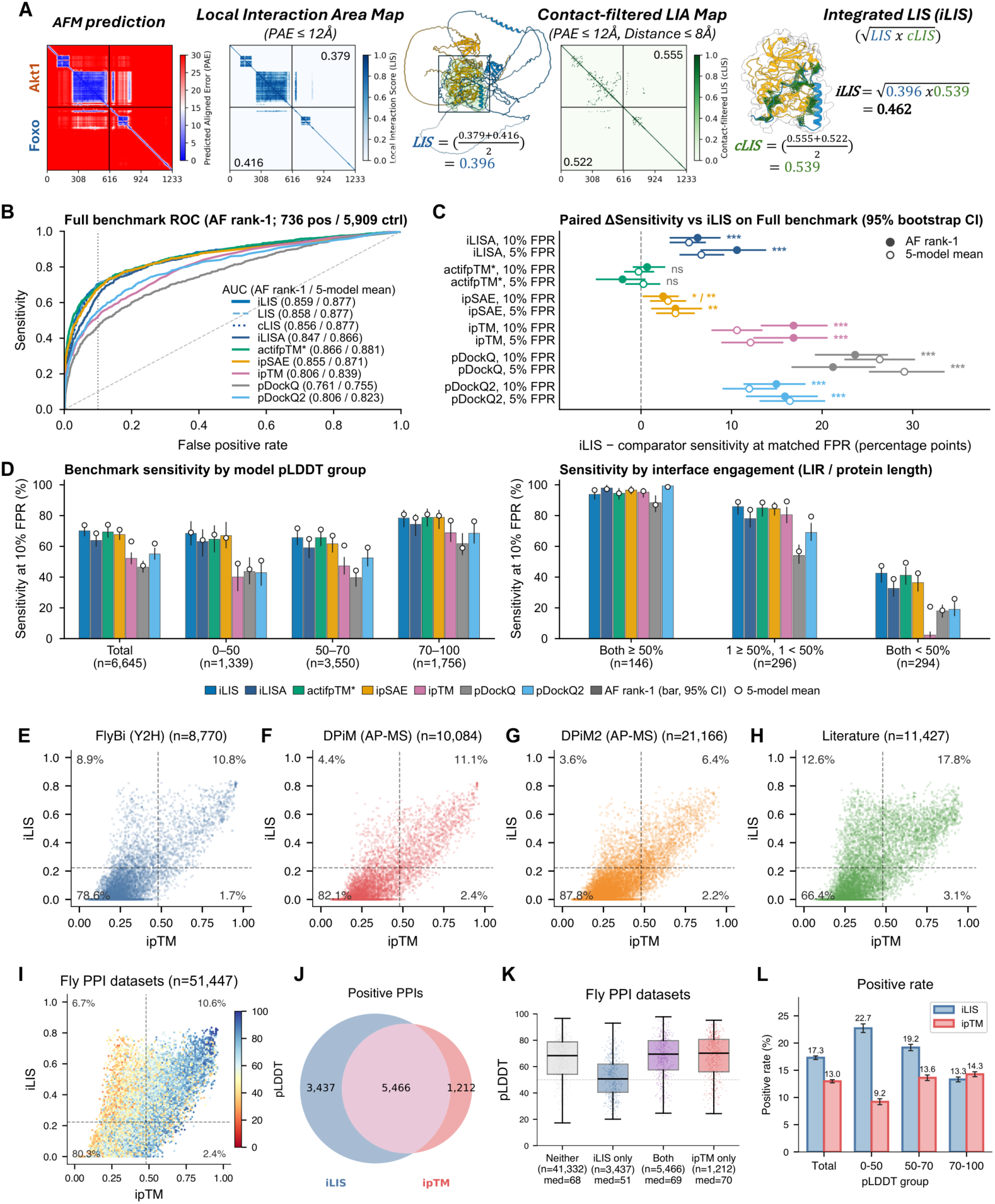
iLIS benchmarking and evaluation against experimental PPI datasets. **(A)** Schematic of iLIS computation (Akt1-Foxo): Local Interaction Area (LIA; PAE ≤ 12 Å) yields LIS (domain confidence) and cLIS (contact confidence, Cβ-Cβ ≤ 8 Å), combined as iLIS = √(LIS × cLIS). **(B)** Full-benchmark AF rank-1 ROC curves (736 positives, 5,909 controls) across nine metrics (legend: AF rank-1 / five-model mean AUC; dotted line, 10% FPR). **(C)** Paired sensitivity differences (iLIS minus comparator, percentage points; 95% stratified bootstrap CIs, 2,000 replicates) at matched 10% and 5% FPR on the full benchmark for AF rank-1 (filled) and five-model mean (open; two-sided paired bootstrap *P*: * < 0.05, ** < 0.01, *** < 0.001, ns not significant; symbols before and after a slash, AF rank-1 and five-model mean). **(D)** Sensitivity at matched 10% FPR by model pLDDT (left: Total, n = 6,645; 0 to 50, n = 1,339; 50 to 70, n = 3,550; 70 to 100, n = 1,756; **Table S3**) and per-chain interface engagement (LIR / length; right: both ≥ 50%, n = 146; one < 50%, n = 296; both < 50%, n = 294; **Table S4**, sheet E3; bars and whiskers, AF rank-1 with 95% CIs; open circles, five-model means). **(E-H)** iLIS versus ipTM (for each metric, the highest value among the five models of a pair; n, records with both scores) for (E) FlyBi Y2H (n = 8,770), (F) DPiM AP-MS (n = 10,084), (G) DPiM2 AP-MS (n = 21,166), and (H) curated literature (n = 11,427); dashed lines, 10% FPR cutoffs (iLIS ≥ 0.223, ipTM ≥ 0.48); quadrant percentages shown. **(I)** Combined scatter across **E-H** (n = 51,447), colored by model pLDDT. **(J)** Overlap of iLIS- and ipTM-positive PPIs (3,437 iLIS-only). **(K)** Model pLDDT boxplots (center line, median; box, interquartile range [IQR]; whiskers, 1.5×IQR) for pairs called by neither (n = 41,332, median 68), iLIS only (n = 3,437, 51), both (n = 5,466, 69), or ipTM only (n = 1,212, 70). **(L)** Positive call rate by pLDDT subgroup for iLIS versus ipTM across **E-H** (0 to 50, n = 10,785; 50 to 70, n = 17,760; 70 to 100, n = 22,902; control FPRs in **Figure S4B**; error bars, Wilson 95% confidence intervals).

We benchmarked iLIS against multiple confidence metrics using 736 positive PPIs (from yeast, fly, and human Y2H PRS and structure-validated interactions from the Eukaryotic Linear Motif [ELM] database; Kumar et al., 2024) and 5,909 controls (Random Reference Sets [RRS], GFP controls, Wingless [Wg] subcellular compartment controls, and shuffled pairs; see Methods for confidence metrics and benchmark dataset, and **Table S1** for the full pair-level dataset). All pairs were modeled from full-length sequences rather than isolated interacting domains, matching how predictions are generated at proteome scale where domain boundaries are rarely known in advance. As a consequence, most models contain substantial non-interacting sequence, including regions of low pLDDT (predicted local distance difference test, AlphaFold’s per-residue confidence score, 0 to 100) unrelated to the interface itself. High pLDDT generally reflects well-folded regions, whereas low pLDDT indicates low prediction confidence, which often coincides with intrinsic disorder but is not equivalent to it (Tunyasuvunakool et al., 2021). Two other local confidence metrics, ipSAE (Dunbrack, 2025) and actifpTM (Varga et al., 2025; computed post hoc from the saved outputs and denoted actifpTM*; Methods), are included here for comparison alongside the established global (ipTM, Model Confidence, and the mean inter-chain PAE, ifPAE [interface PAE]) and distance-defined contact-based metrics (pDockQ, pDockQ2, ifPAE_d8), as well as iLISA, an area-weighted variant of iLIS (Methods; **Table S2**).

Spearman correlation analysis revealed distinct clustering among metrics, with the local metrics correlating most strongly with positive PPIs (**Figure S3A-B**). In receiver operating characteristic (ROC) analysis of the full benchmark, the local confidence metrics (iLIS, ipSAE, and actifpTM*) exceeded the global metric ipTM and the contact-based metrics pDockQ and pDockQ2 (**Figure 1B**; **Table S3**). At matched false positive rates (FPR) of 10% and 5%, iLIS recovered 7 to 17 percentage points more reference interactions than ipTM and 11 to 29 more than pDockQ on the calibration set and the full benchmark, and more on the held-out motif set (**Figures 1C**, **S4A**). Among the three local metrics, the differences were smaller: iLIS and actifpTM* did not differ significantly at these rates, iLIS exceeded ipSAE by at most 4 points on the calibration set and the full benchmark (7 points at 5% FPR on the held-out motif set), and actifpTM* recovered 6 to 17 points more interactions at the 1% FPR tier on single models, a difference in sensitivity that disappeared with five-model averaging (paired comparisons in **Table S4**).

Because full-length modeling means each pair carries a different proportion of confidently versus poorly predicted sequence, we stratified performance by the model’s overall pLDDT (0 to 50, low confidence; 50 to 70, intermediate; 70 to 100, high confidence). The local confidence metrics consistently exceeded global metrics such as ipTM and other widely used metrics such as pDockQ across all three pLDDT subgroups (**Figure 1D**, **left**; **Figure S3C-D**), with particularly clear separation for low-pLDDT pairs, where a metric that isolates the interface from surrounding low-confidence sequence matters most (**Figure S3E-H**; bootstrap benchmarking results for all metrics in **Table S3**).

Because global metrics average across entire chains, this separation tracked the fraction of each protein participating in the interface (Local Interaction Residues [LIR], the residues in the confident interaction domain, divided by protein length; **Figure 1D**, **right**). When the confident interaction domain covered at least half of both chains, all metrics performed comparably (88 to 99% sensitivity at 10% FPR), and when it covered at least half of only one chain, ipTM remained close to iLIS (80.5% versus 85.8%). In contrast, when the interface spanned less than half of both chains, non-interacting sequence diluted ipTM to 2.3% sensitivity, whereas iLIS retained 42.5% (**Figure 1D**, **right**; **Table S4**). Within this group, iLIS recovered 80.8% of the positives in which both chains engaged 20 to 50% of their length (n = 26) and 22.1% of those in which both engaged less than 20% (n = 140), compared with 4.9% and 0.0% for ipTM on AF rank-1 models (the model ranked first by AlphaFold-Multimer; 65.4% and 0.7% for ipTM with five-model averaging; **Figure S3I-J**).

As an independent test, we repeated the comparison on complexes deposited in the Protein Data Bank (PDB) after the AFM training cutoff (338 complexes against 1,257 GFP or Wg controls; **Figure S5**). On these mostly compact, confidently modeled constructs (median pLDDT 89), iLIS (area under the ROC curve [ROC-AUC] 0.895), ipSAE (0.898), actifpTM* (0.892), and ipTM (0.888) did not differ significantly, in ROC-AUC or in precision-recall AUC (0.838 for iLIS, 95% confidence interval [CI] 0.806 to 0.869), and iLIS retained its discrimination among complexes with < 30% sequence identity to any pre-cutoff PDB chain (ROC-AUC 0.925, n = 38) and among the 109 complexes for which no single pre-cutoff entry contained homologs of both chains (0.868, 95% CI 0.826 to 0.910). The advantage of local metrics over ipTM is therefore small on compact complexes. No difference was seen either when 140 of these complexes were re-predicted from full-length sequences (ROC-AUC 0.841 for both iLIS and ipTM; **Figure S5D, E**), whereas in the reference benchmark the advantage was largest for pairs in which the interface covered less than half of both chains (**Figure 1D**, **right**).

To test whether iLIS also tracks model accuracy rather than only classification, we compared it with each model’s DockQ against the deposited experimental structure across 402 unique post-cutoff complexes (the 338 construct models plus 64 complexes modeled only at full length; Methods): iLIS correlated with DockQ (Spearman ρ = 0.76), including among the 49 complexes with < 30% identity to any pre-cutoff PDB chain (ρ = 0.70, P = 2 × 10^-8^; **Figure S3K**; **Table S5**).

To enable downstream PPI classification at large scale, we calibrated thresholds on the Y2H-derived PRS from the yeast, fly, and human proteome-scale screens as positives and on the Y2H-derived RRS, GFP, and Wingless control sets as negatives (n = 357 positives and 1,255 negatives with complete scores across all metrics; the ELM interactions and shuffled pairs were not used for calibration and served as a test set; Methods). Thresholds were computed across a range of FPRs and are provided for every metric in **Table S6**. Tightening the cutoff carries a steep sensitivity cost: for iLIS, sensitivity falls from 73% at 10% FPR (≥ 0.223) to 67% at 5% (≥ 0.339), 38% at 1% (≥ 0.551), and 15% at a nominal 0.1% (≥ 0.662). We therefore adopted iLIS ≥ 0.223 (10% FPR) for all interaction calls in this study.

Applied without recalibration, the iLIS ≥ 0.223 cutoff retained its sensitivity on held-out ELM interactions (73.9%, 95% CI 69.0 to 78.4%) and post-cutoff PDB complexes (77.5%). However, the achieved FPR varied with the control population: 8.1% for post-cutoff PDB controls (95% CI 6.7 to 9.8%) and 15.1% for held-out controls (14.1 to 16.2%), where shuffled motif pairs (18.1%) and pairs with model pLDDT below 50 (20.3%) pass most frequently (**Figures S4B, S5F-H**; **Table S6**). The nominal 10% therefore describes the calibration controls and is not a fixed property of the cutoff.

At this threshold, iLIS rescued 47 PRS interactions below the matched-FPR ipTM threshold (≥ 0.48; **Figure S1C-D**), including examples shown in **Figure S1A**; across the full benchmark, 83% of iLIS-only positives engage less than half of both chains (**Figure S1E**) and their interfaces are compact (median 46 contact residues versus 65 for positives called by both metrics; **Table S4**). When positives and controls were both counted among discordant calls, iLIS-only pairs (relative to ipTM) were enriched 3.9-fold for reference interactions in the calibration set (39/357 vs. 35/1,255) and 4.5-fold in the held-out set (93/330 vs. 292/4,620), whereas ipTM-only and pDockQ-only calls were depleted (likelihood ratios 0.45 to 0.70 and 0.24 to 0.57; **Table S4**; representative cases in **Figure S6**). True and false iLIS-only calls share low overall pLDDT and differ in contact area and in pLDDT at the contact residues (**Table S4**). Discordance between iLIS and the other local metrics involved few pairs (**Figure S7A, B**), and neither direction was enriched for true interactions (for iLIS versus ipSAE on the calibration set, 3 positives and 20 controls were called only by iLIS and 6 positives and 21 controls only by ipSAE; **Table S4**).

Agreement among the four metrics (iLIS, ipSAE, actifpTM*, ipTM) depended on the stringency of the cutoff and on the population scored. Among the Y2H benchmark positives, the four metrics agreed on 61.6% of the true interactions at their deployed 10% FPR thresholds, a concordant core that narrowed to 22.1% at 1% FPR as each metric retained a different subset (**Figure S7A**). Measured on the same denominator as for the atlas (all pairs called positive by any of the four metrics), benchmark agreement was 80.6% at 10% FPR and 38.0% at 1% FPR. Across the full atlas described below, the metrics agreed less: of all pairs called positive by any metric, only 35.4% were called by all four at 10% FPR and 12.9% at 1% FPR, because the atlas contains many borderline predictions where the metrics diverge (**Figure S7C**).

Taken together, iLIS performs on par with ipSAE and actifpTM* and exceeds global and contact-based metrics on the full-length reference benchmark. We use it as the primary metric because its contact term identifies the residues in direct contact, on which the interface analyses below depend, and we report the other metrics for every prediction in the database. Because no metric is uniformly best, the database also gives a composite tier score that sums, over five metrics, the FPR tiers a pair passes, as an alternative for users who prefer not to choose a single metric. At the 10% tier, the composite score performs like iLIS, both on the calibration set (73.9% sensitivity at 9.9% FPR for a composite score of at least 3 of 15, versus 72.8% at 10.0% for iLIS), and on held-out pairs (72.4% at 15.0%, versus 73.9% at 15.1%). Calling a pair when any of four metrics passes recovers about as many reference interactions as relaxing the iLIS cutoff to the same number of false positives, so the other metrics offer an alternative ranking and do not add interactions beyond a more permissive cutoff (**Table S4**). The analyses below use iLIS calls at the 10% FPR cutoff, while the companion metrics and the composite score allow borderline calls to be examined.

### Benchmarking iLIS across experimental and predicted PPI sets

We next evaluated predictions against large-scale *Drosophila* experimental PPI datasets, including the FlyBi Y2H screen (Tang et al., 2023) and the DPiM/DPiM2 AP-MS interactomes (Guruharsha et al., 2011; Bhat et al., 2024). We recovered approximately 20% of FlyBi, 16% of DPiM, and 10% of DPiM2 interactions as positive (iLIS ≥ 0.223; all reported records, **Figure 1E-G**). We further assessed recovery of a large set of literature-reported *Drosophila* PPIs sourced from FlyBase (Öztürk-Çolak et al., 2024), BioGRID (Oughtred et al., 2021), and MIST (Hu et al., 2018). Of these curated PPIs, ∼30% were positively predicted (**Figure 1H**). This higher recovery likely reflects that literature-curated interactions are typically validated through focused biochemical studies.

Relative to atlas pairs among the same proteins that a dataset did not report, reported pairs were called positive more often for FlyBi, DPiM, and the curated literature (Mantel-Haenszel odds ratios 1.57, 1.63, and 1.92), but not for DPiM2 (0.67). Against randomly paired fly proteins, which were called at 8.8% (random reference set) and 17.5% (shuffled reference pairs), the recovery of the three proteome-scale screens, counted on unique heterodimer pairs (17.4%, 15.3%, and 10.0%), did not exceed the higher of the two control rates (**Table S4**). The recovery rates of these screens therefore do not by themselves indicate enrichment above background: for FlyBi and DPiM, enrichment rests on the comparison within the screened proteins, and for DPiM2, recovery was lower than among the non-reported pairs of the same proteins.

Across all datasets, iLIS called reported pairs that scored below the ipTM threshold (**Figure 1E-I**), 3,437 pairs in total (**Figure 1J**), and these iLIS-only pairs had lower pLDDT than pairs called by both metrics (**Figure 1I**, **K**). Because these datasets contain no verified negatives, the additional calls raise the positive call rate but do not by themselves measure precision; the error rate of iLIS-only calls is estimated on the reference benchmark above. The difference in call rate between iLIS and ipTM was largest at low pLDDT (0 to 50), where iLIS called 22.7% of reported pairs compared to 9.2% for ipTM (**Figure 1L**) and where the control FPR of iLIS is also highest (20.3% versus 11.7% for ipTM on held-out controls; **Figure S4B**).

The positive prediction rate tended to increase with the directness of experimental evidence, from ∼10 to 20% for proteome-scale PPI screens (FlyBi, DPiM, and DPiM2) to 68 to 81% for direct-binding and structural methods (enzyme-linked immunosorbent assay, surface plasmon resonance, X-ray crystallography, and isothermal titration calorimetry; **Figure S8A**). It also increased with the depth and breadth of supporting evidence: from 17.5% for pairs backed by a single experimental record to 75.5% for those backed by ten or more independent records (**Figure S8B**), and from 19% for pairs reported by a single method to 65% for those reported by four or more distinct methods (**Figure S8C**). A pairwise call does not by itself distinguish direct contact from co-complex membership, because subunits of one complex that do not touch can still be docked when predicted as a pair (see the analysis of higher-order assemblies below).

We extended this analysis to three recent computational PPI prediction datasets. On a *Drosophila* AlphaFold-Multimer set from Peng and Zhao (2026; 27,711 predictions, 27,683 unique gene pairs after our mapping; classified by pDockQ), 5,601 pairs were called by both iLIS (≥ 0.223) and pDockQ (≥ 0.23), 2,505 by pDockQ only, and 1,070 by iLIS only; all of the iLIS-only pairs fall in the class that Peng and Zhao labeled “possible” (**Figure S9A,B**).

Comparing FlyPredictome against Peng and Zhao and against the AlphaFold Database (Han et al., 2026) showed limited direct overlap (**Figure S9C**). To test whether this reflects the largely non-overlapping candidate sets each resource screened rather than disagreement among the metrics, we rescored the complete *Drosophila* heterodimer set of Han et al. (138,938 models, including those below their high-confidence threshold). Only 29,706 of its gene pairs were also predicted in FlyPredictome; for the 29,685 shared pairs with model outputs, matched single models from the two resources agreed in 92.3% of calls at iLIS ≥ 0.223 (Spearman ρ = 0.65), most of them pairs negative in both (24,050); of the 5,635 pairs positive in either resource, 3,339 were positive in both. In addition, 95.8% of the shared AlphaFold Database high-confidence pairs (865 of 903) are positive in FlyPredictome (**Figure S9D-F**). The two resources therefore differ mainly in the candidate pairs screened and the threshold applied.

Applied to the human predictome of Schmid et al. (2025; 1,614,047 pairs), iLIS recovered 99.5% (16,479 of 16,557) of their SPOC classifier high-confidence calls while nominating 210,651 further positives (227,130 in total; **Figure S9G**; **Table S7**). The two resources also overlapped substantially across species: 47,783 of the 73,899 fly positives whose human ortholog pair Schmid and colleagues had modeled (∼65%) were positive in the human predictome as well, largely independent of how confidently the ortholog was assigned (**Figure S9H**); the corresponding rate for fly pairs below the cutoff was not computed, so this overlap is descriptive. Those 47,783 pairs are about a third of all 144,779 fly positives, with most of the remainder falling outside the pairs Schmid et al. screened rather than reflecting scoring disagreement (**Figure S9I**); in the reverse direction, 54,227 of 225,749 Schmid et al. human positives likewise had a fly-positive ortholog (**Figure S9J**). Together, these analyses show that iLIS calls are concordant with independently generated prediction datasets on the pairs they share. Because these resources also rely on AlphaFold-Multimer, the concordance does not establish that the shared calls are correct.

### Predicted structural models across *Drosophila* pathways

Beyond metric benchmarking and recovery analysis, we asked whether these structural models could predict the direct physical interactions that underlie established signaling pathways in which the pathway components are well defined genetically but the structural basis of many individual binding events remains unknown. As a representative example, **Figure 2** illustrates the predicted structural complexes along the Epidermal Growth Factor Receptor (EGFR)/Ras-MAPK cascade, one of the best-characterized receptor tyrosine kinase pathways in *Drosophila* (reviewed in Sopko and Perrimon, 2013; Shilo, 2014). Our predictions provide structural models for direct PPIs throughout the cascade, from ligand regulation (Argos-Spitz) and receptor activation (Egfr-Spitz) to adaptor recruitment (Egfr-Shc), phosphatase recruitment (Corkscrew-Dos), scaffold assembly (KSR-Dsor1), and the terminal MAPK relay (Rolled-Dsor1).

**Figure 2.**
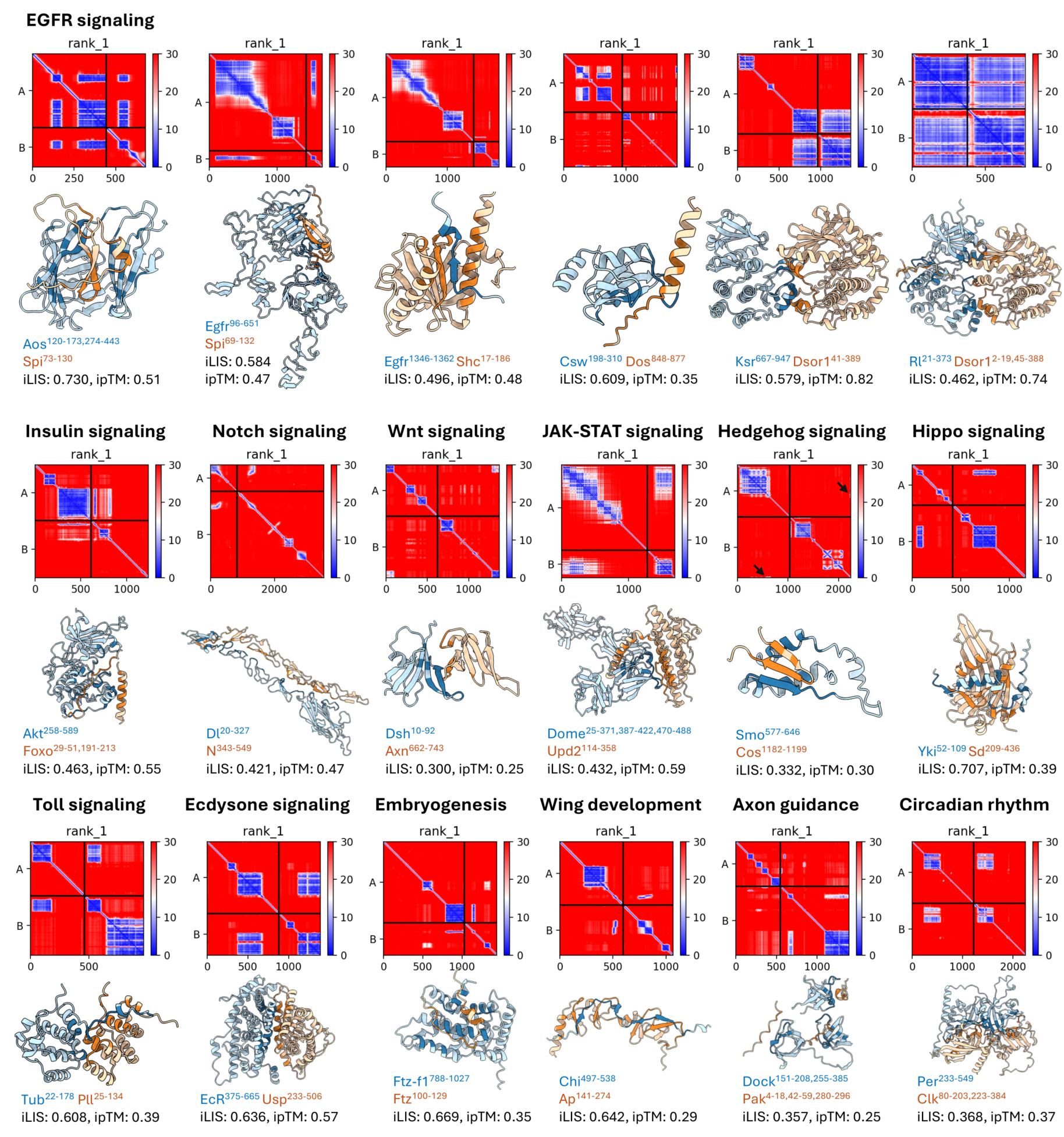
Predicted structural models of major signaling pathway PPIs. AFM-predicted structural complexes along the EGFR/Ras-MAPK cascade (top row) and across 12 additional signaling contexts (middle and bottom rows). For each pair, the Predicted Aligned Error (PAE) heatmap (top) and predicted 3D structure (bottom) are shown. In the PAE heatmaps, each cell represents the expected positional error (in Å) between two residues; low values (dark blue) indicate high-confidence relative positioning, while off-diagonal blue blocks between the two chains mark the predicted interaction interface. In the structures, chain A is colored in blue and chain B in orange, indicating Local Interaction Residue (LIR), with darker shading indicating contact LIR (cLIR). Residue ranges of the interacting domains and the iLIS and ipTM values are indicated below each structure; only AF rank-1 predictions are shown. Black arrows highlight small interaction interfaces (in Smo-Cos). Some complexes, such as Dsh-Axn (iLIS = 0.300, ipTM = 0.25) and Smo-Cos (iLIS = 0.332, ipTM = 0.30), have low ipTM scores despite passing the 10% FPR iLIS cutoff (0.223; both fall below the 5% FPR cutoff of 0.339).

These predictions extend beyond a single signaling cascade. **Figure 2** further catalogs predicted complexes across 12 additional biological contexts spanning major signaling pathways, developmental processes, and behavioral programs. Notably, Chip binds Apterous LIM domains (Morcillo et al., 1997), and GST pulldown with deletion constructs mapped this interaction to the C-terminal LIM Interaction Domain (LID) of Chip (van Meyel et al., 1999; Torigoi et al., 2000); our predicted Chip-Apterous interface coincides with this experimentally defined region, illustrating that iLIS can recover interfaces mapped through biochemical assays.

Because confidence alone does not distinguish compact motif docking from extensive domain-domain binding, we also evaluated interface extent using the area-weighted score iLISA (iLIS multiplied by the interaction area iLIA; Methods) across 25 established *Drosophila* complexes and the EGFR/Ras-MAPK cascade (**Figure S10**). For large-interface complexes with modest iLIS, such as aPKC-l(2)gl (rank 54 to 1 of 1,579 modeled partners) and arm-alpha-Cat (rank 7 to 1 of 760), weighting by interface area elevated the canonical partner to the top rank, whereas compact motif interactions such as key-Atg8a (rank 1 to 108) were prioritized by iLIS but demoted by iLISA (**Figure S10A**). Similarly, across the EGFR/Ras-MAPK cascade, iLISA surfaced large kinase and scaffold docking interfaces, including Raf-Dsor1 and Dsor1-Rl (**Figure S10B-G**), enabling prioritization of either compact motif-mediated contacts with iLIS or extensive domain-domain interfaces with iLISA depending on the biological focus.

### Missense alleles are enriched at predicted interfaces

Next, to assess whether the predicted interfaces are functionally important, we examined whether missense alleles (point mutations associated with phenotypic effects identified through genetic studies) are preferentially associated with predicted interaction sites. Missense mutations were compiled from FlyBase allele descriptions and genetic interaction records, and each was mapped to the corresponding residue in AFM-predicted structural models (see Methods). After filtering for validated amino acid identity at the mapped position, removing synonymous entries, and restricting to proteins whose interface profiles were built from *Drosophila* predictions, 2,504 unique mutations across 861 proteins were retained for analysis. For each residue, we asked two questions across all of a protein’s modeled partners: how often it lies within a partner’s interaction domain, and how often it sits at the direct contact interface. These define two residue-level frequencies: the LIR percentage, the fraction of partner models in which the residue falls within the confident interaction domain (PAE ≤ 12 Å), and the contact LIR (cLIR) percentage, which further restricts to residues in direct intermolecular contact (Cβ-Cβ ≤ 8 Å and PAE ≤ 12 Å).

We tested whether mutation rate increases with LIR percentage across these 861 proteins. The mutation rate increased monotonically from 0.10% for residues never in any interaction domain to 0.71% for the highest quartile, a 6.9-fold difference before any adjustment (**Figure 3A**). Among residues within an interaction domain, there was additional enrichment at the direct contact interface. The mutation rate increased from 0.42% to 0.87%, a 2.1-fold enrichment (**Figure 3B**). Within-gene permutation tests showed that both trends exceed what differences between genes in the number of mutations would produce (P < 10^-5^ for both; **Figure S11A-B**).

**Figure 3.**
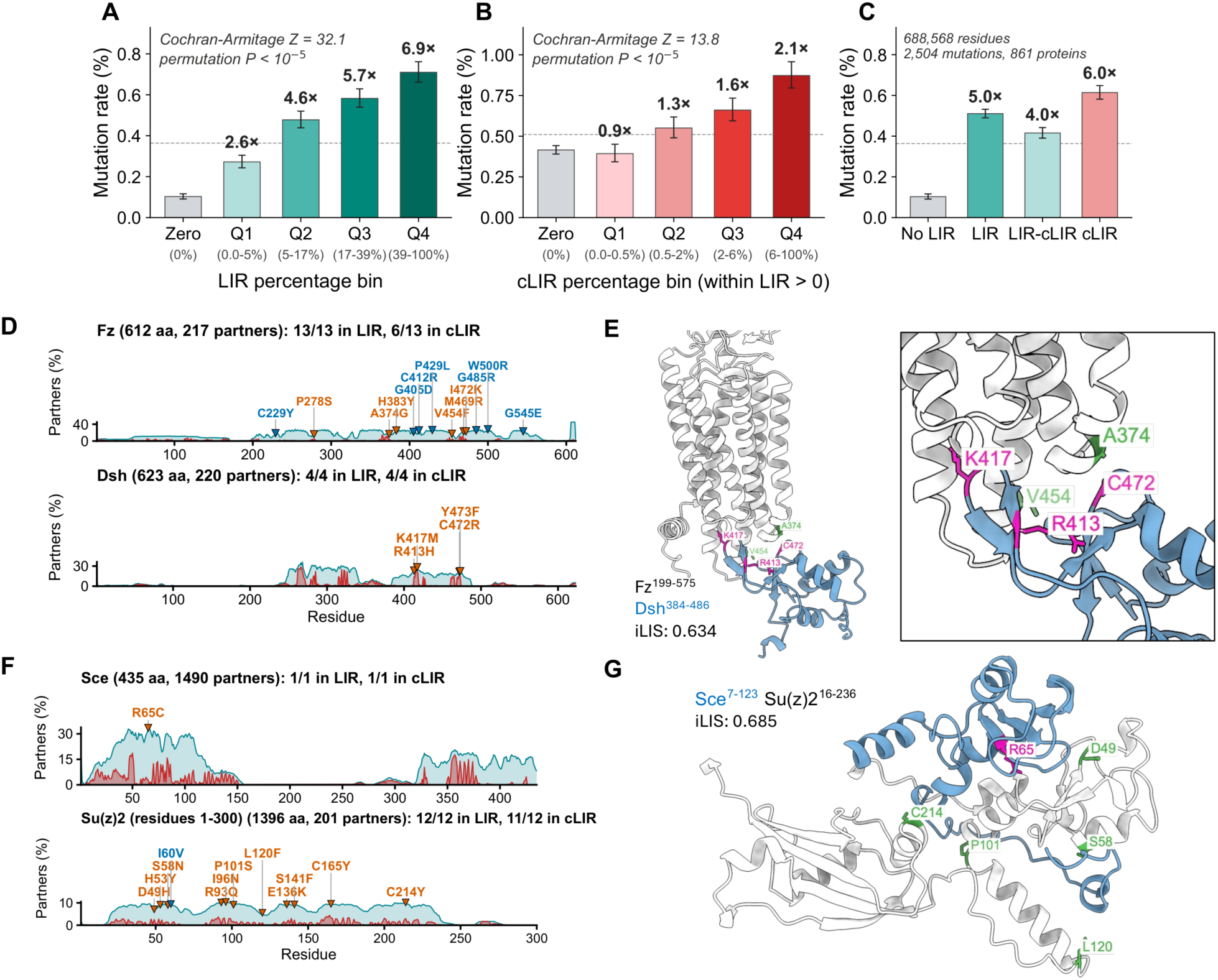
Missense alleles are enriched at predicted interaction interfaces. **(A)** Mutation rate across LIR frequency bins for 688,568 residues in 861 proteins (2,504 missense mutations). Residues are binned into Zero (never in any interaction domain) and quartiles Q1-Q4 of the non-zero distribution. Bars show the mutation rate (mutated residues / residues in the bin) and error bars its Wilson 95% confidence interval; fold-enrichment relative to Zero is indicated above each bar. Bin sizes and counts are provided in **Table S4** (sheet S). Cochran-Armitage trend test Z = 32.1 (permutation P < 10^-5^). Dashed lines in panels **A**-**C** indicate the average mutation rate. **(B)** Mutation rate across cLIR frequency bins for residues within predicted interaction domains (LIR > 0). Cochran-Armitage trend test Z = 13.8 (permutation P < 10^-5^). **(C)** Mutation rate (bars, Wilson 95% confidence intervals) across residue categories: No LIR (not in any interaction domain), LIR (in interaction domain), LIR−cLIR (in interaction domain but not at contact), and cLIR (direct contact). All three interaction categories show elevated mutation rates relative to No LIR; for adjustment by prediction confidence, see **Figure S12**. **(D)** LIR (teal) and cLIR (salmon) frequency profiles for Frizzled (Fz; top) and Dishevelled (Dsh; bottom). Each profile shows the fraction of modeled partners in which a given residue is part of the interaction domain (LIR) or at direct contact (cLIR). Vermilion and blue triangles indicate mapped missense mutations at direct-contact residues and at interaction-domain residues without direct contact, respectively. For Fz, all 13 mutations fall within the interaction domain. For Dsh, all 4 mutations fall at direct contact residues. **(E)** Predicted wild-type Fz-Dsh complex (iLIS = 0.634). Mutation positions in Dsh (magenta) and Fz (green) are highlighted at the binding interface. **(F)** LIR/cLIR profiles for Sce (top) and Su(z)2 (bottom; residues 1 to 300), with mutations marked as in **D**. The single Sce mutation (R65C) falls at a direct contact residue. Of 12 mapped Su(z)2 mutations, 11 fall at direct contact residues. **(G)** Predicted wild-type Sce-Su(z)2 complex (iLIS = 0.685). Mutation positions in Sce (magenta) and Su(z)2 (green) are highlighted at the binding interface.

To ask whether the enrichment extends beyond direct contacts, we separated domain residues into those making direct contact (cLIR) and those in the domain but not at the contact interface (LIR−cLIR). Compared to non-interacting residues (0.10%), all interaction-domain residues showed elevated mutation rates (LIR: 0.51%; **Figure 3C**). Within these, domain residues lacking direct contact (LIR−cLIR: 0.42%) were intermediate between non-interacting and direct contact residues (cLIR: 0.61%). Even among domain-only residues (LIR > 0, cLIR = 0), mutation rate increased monotonically with LIR percentage (3.1 to 5.3× enrichment; permutation P = 2 × 10^-5^; **Figure S11C-D**). This domain-level trend is not adjusted for prediction confidence, which we examine below. Overall, 90% of mutations fell within a predicted interaction domain and 51% at a direct contact residue, compared with 79% and 43% expected if each mutation were placed at random within its own protein (64% and 30% of all residues pooled; **Table S4**).

The enrichment at contact residues is not explained by residue burial. Interaction-domain residues are more often buried than background residues: 29% of cLIR residues and 56% of domain-only residues are buried (relative solvent accessibility [RSA] < 0.25), compared with 14% of background residues (**Figure S11E-F**). To test whether this difference in burial accounts for the enrichment, we compared residues within the same burial class: mutation enrichment persists within each class, with cLIR residues enriched 5.0-fold among surface and 4.8-fold among buried residues relative to background of the same burial class (**Figure S11G**), and cLIR remained a significant predictor of mutation status after adjusting for RSA (odds ratio 2.5 for contact versus all other residues; P < 10^-100^, treating residues as independent).

Evolutionarily conserved residues tend to produce phenotypes when mutated, raising the possibility that the observed enrichment reflects conservation rather than interface-specific constraint. To test this, we controlled for evolutionary conservation using PhyloP scores (Pollard et al., 2010) from the 27-way insect species alignment (23 *Drosophila* species and 4 outgroup insects) available through the UCSC Genome Browser (Kent et al., 2002). Interface residues were indeed more conserved than non-interface residues (mean PhyloP: non-interface 1.84, interface 2.62; **Figure S11H**). However, when we stratified residues into four equal groups by conservation level, the interface enrichment persisted across all conservation levels (3.5 to 7.1×, with the highest enrichment in intermediate-conservation residues rather than the most conserved; **Figure S11I**). Logistic regression confirmed that cLIR remained a highly significant predictor of mutation status after adjusting for PhyloP (P = 3 × 10^-59^), with conservation accounting for only an 8.6% attenuation of the interface effect (**Figure S11J**).

Enrichment at contact residues was observed in each phenotype-severity class: missense alleles categorized as lethal (52.3% at contact, versus 43.3% expected within the same proteins; P = 9 × 10^-7^), semi-lethal (62.3% versus 40.1%; P = 9 × 10^-4^), and viable/mild (47.8% versus 40.7%; P = 4 × 10^-4^) were each over-represented, with no significant gradient across classes (**Figure S11K**, in which the dashed lines show the pooled residue background).

Cross-referencing predicted interfaces with curated post-translational modification sites (UniProt/Swiss-Prot) further showed that phosphosites are specifically enriched at direct-contact residues (37.1%) relative to other Ser/Thr/Tyr residues without a curated phosphosite in proteins carrying a curated modification site (26.4%; odds ratio 1.65, P = 5 × 10^-9^; **Figure S11L**), consistent with phosphorylation at binding contacts or with phosphoacceptor sites engaged by predicted kinase partners.

Finally, we asked whether the enrichment reflects prediction confidence and the depth of the multiple sequence alignment (MSA) rather than interfaces themselves (**Figure S12**). In a joint logistic regression that included residue pLDDT, RSA, PhyloP, LIR percentage (as a measure of local inter-chain confidence), and gene fixed effects, direct-contact frequency remained associated with mutation (odds ratio 1.19 per standard deviation of cLIR, 95% CI 1.16 to 1.22; 1.20 without adjustment). The estimate was unchanged when MSA depth was added and was 1.18 when all structural covariates were derived from the same MSA, and contact residues were enriched over residues outside any interaction domain within every quartile of pLDDT and of MSA depth (2.8- to 4.5-fold). In contrast, the graded increase with LIR percentage was largely explained by prediction confidence (odds ratio 1.05, 95% CI 0.98 to 1.14, in the same model; 1.70 without adjustment). The enrichment at direct-contact residues therefore persists after these adjustments, whereas the graded domain-level trend is largely accounted for by the covariates, chiefly prediction confidence.

### Allele enrichment in well-characterized signaling proteins

To visualize this enrichment in individual proteins, we first examined Wnt/planar cell polarity (PCP) signaling, where the receptor Frizzled (Fz) couples to Dishevelled (Dsh) via the Dsh DEP domain. Among the 13 mapped mutations of Fz, 6 fell at direct contact residues; for Dsh, all 4 mapped mutations were at direct contact residues (**Figure 3D**). When we mapped these mutations onto the predicted Fz-Dsh complex, 2 Fz mutations (*fz[24]*: A374G; *fz[GL31]*: V454F) and 3 Dsh mutations (*dsh[1]*: K417M; *dsh[A3]*: R413H; *dsh[A21]*: C472R) mapped to the binding interface (**Figure 3E**). The three Dsh mutations all reside in the DEP domain and have been shown to specifically disrupt PCP signaling (Axelrod et al., 1998; Boutros et al., 1998; Penton et al., 2002), consistent with selective disruption of the Fz-Dsh interface. A recent DVL2-FZD4 cryo-EM structure (PDB: 8WM9; Qian et al., 2024), published after the AFM training data cutoff, closely matches our predicted Fz-Dsh complex (**Figure S13A**), supporting the predicted interface to which these PCP-specific alleles map.

As a second example, we examined the Polycomb Repressive Complex 1 (PRC1) ubiquitylation module components Sce (Sex combs extra/dRING, the catalytic E3 ligase) and Su(z)2, a paralog of the canonical PRC1 subunit Psc. Of the 12 mapped mutations in Su(z)2, 11 fell at direct contact residues across predicted binding partners, and 5 map specifically to the predicted Sce-Su(z)2 interface. The single Sce mutation (R65C) also maps to this interface (**Figures 3F**, **3G**). The predicted RING-RING interface matches the known Ring1B-BMI1 crystal structures (PDB: 2CKL, Buchwald et al., 2006; PDB: 2H0D, Li et al., 2006) (**Figure S13B-C**). The prediction also suggests contacts extending beyond the crystallized RING domain into the RAWUL domain of Su(z)2, where the C214Y mutation maps (Nguyen et al., 2017).

This pattern extends beyond these two examples: LIR/cLIR profiles for 10 additional genes show alleles clustering at predicted interaction hotspots (**Figure S14**), and structural examination of 16 additional complexes across diverse signaling and regulatory contexts illustrates missense alleles mapping to or near the predicted binding interface in each of these examples (**Figure S15**). Together, these findings indicate that predicted interaction interfaces are enriched for functionally important residues, as independently defined by classical genetics.

### FlyPredictome database and evidence-supported PPI network

With iLIS benchmarked as a confidence metric and predicted interfaces supported by allele enrichment, we scaled our predictions to build FlyPredictome, a database for *in silico* PPI discovery (https://www.flyrnai.org/tools/fly_predictome; **Figure 4A**). The database contains 1.5 million pairwise AFM predictions, including homodimer predictions for nearly all *Drosophila* proteins, and corresponds to 1,108,443 unique heterodimeric protein pairs (∼1.1% of all possible *Drosophila* pairs) assembled from targeted prediction sets: 573,127 from the secreted ligand × receptor extracellular-domain screen, 259,217 from the kinase interactome, 181,014 from other targeted, pathway, and collaboration sets, 79,636 from previously reported interactions and large-scale proteomics datasets, and 15,449 from organelle-targeted sets. Protein pools, pairing rules, and input sequences for every set are described in Methods, and their composition and positive rates are enumerated in **Table S8**. After consolidating overlapping prediction sets and isoforms to unique protein pairs, 144,779 heterodimeric PPIs and 5,470 homodimers were classified as positive (iLIS ≥ 0.223; **Figure 4B**).

**Figure 4.**
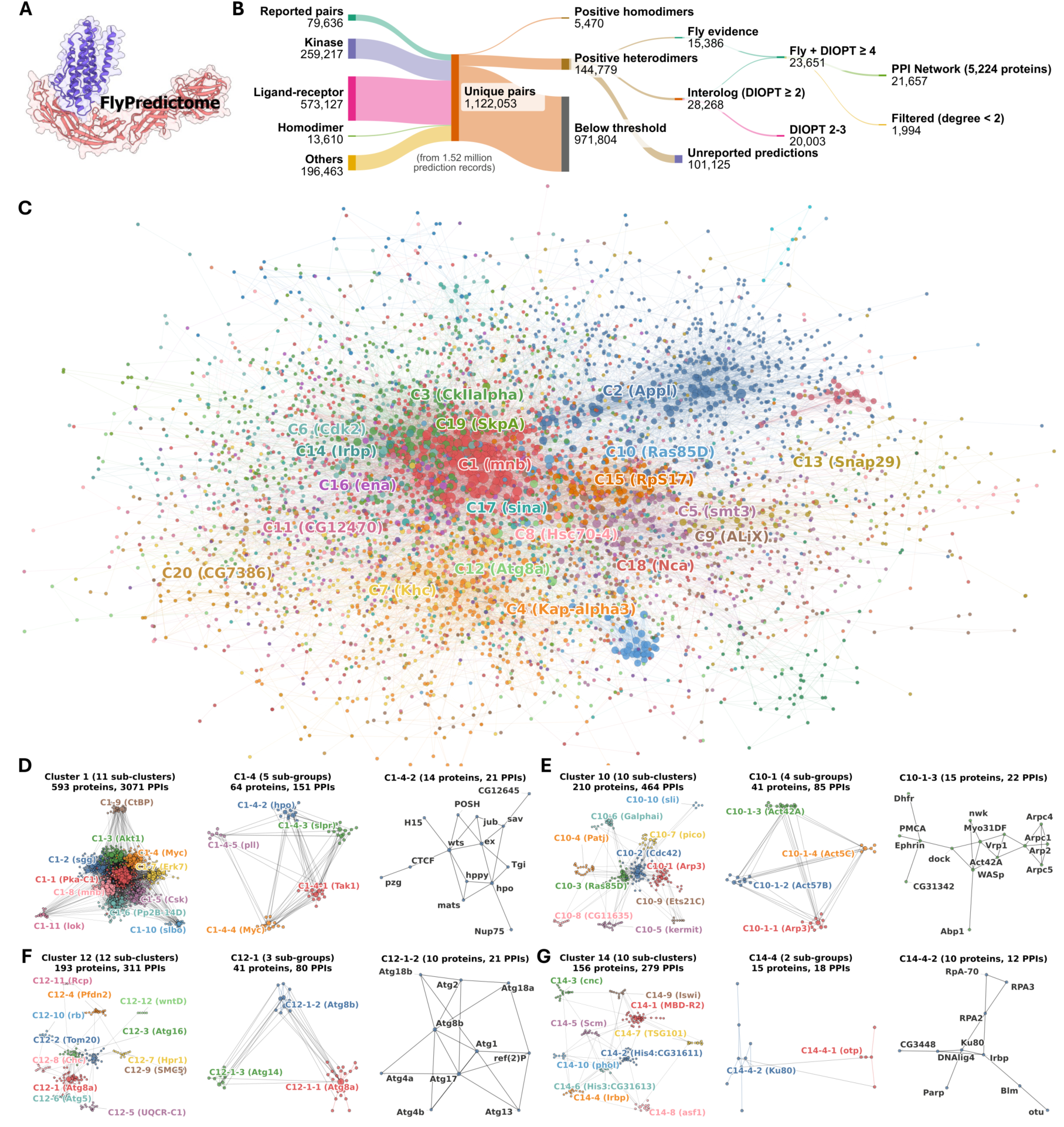
FlyPredictome database and evidence-supported PPI network. **(A)** The FlyPredictome database (https://www.flyrnai.org/tools/fly_predictome). **(B)** Sankey diagram showing the flow from 1,122,053 unique protein pairs (1,108,443 heterodimeric and 13,610 homodimeric, consolidated from 1.52 million AFM prediction records and grouped by source as reported pairs, kinase pairs, ligand-receptor pairs, homodimers, and others; **Table S8**) through iLIS classification (144,779 positive heterodimers and 5,470 positive homodimers), evidence annotation (fly literature, interolog conservation at DIOPT ≥ 2, or unreported), to the final evidence-supported network (fly literature or DIOPT ≥ 4; 5,224 proteins, 21,657 PPIs), shown in **C**. All predictions, including those below threshold, are accessible through FlyPredictome. **(C)** Evidence-supported PPI network colored by cluster assignment. Leiden community detection identified 31 clusters; the 20 largest are annotated with their hub protein. Intra-cluster edges are colored by cluster; inter-cluster edges are gray. An interactive version is available at https://flyark.github.io/FlyPredictome-network. **(D)** Hierarchical decomposition of Cluster 1 (kinases/transcriptional regulators; panels **D-G** are enlarged in **Figure S23**). Left: full cluster (593 proteins, 11 sub-clusters). Center: C1-4 (JNK cascade/growth control; 5 sub-groups). Right: C1-4-2 (14-protein Hippo signaling module). **(E)** Hierarchical decomposition of Cluster 10 (signal transduction/cytoskeleton). Left: full cluster (210 proteins, 10 sub-clusters). Center: C10-1 (Arp2/3 actin nucleation; 4 sub-groups). Right: C10-1-3 (Arp2/3 and actin regulatory module). **(F)** Hierarchical decomposition of Cluster 12 (autophagy). Left: full cluster (193 proteins, 12 sub-clusters). Center: C12-1 (macroautophagy; 3 sub-groups, including C12-1-3, the PI3K/Vps34 nucleation complex). Right: C12-1-2 (autophagosome assembly module). **(G)** Hierarchical decomposition of Cluster 14 (chromatin/DNA repair). Left: full cluster (156 proteins, 10 sub-clusters). Center: C14-4 (NHEJ DNA repair; 2 sub-groups). Right: C14-4-2 (NHEJ ligation complex). Cluster assignments, GO enrichment, network edges, and evidence composition are provided in **Tables S11-S15**.

Because the ligand-receptor screen used truncated receptor extracellular domains (ECDs) whereas the cutoff was calibrated on full-length pairs, we tested the cutoff on ECD-matched negative controls: each of 421 receptor ECD isoforms paired with GFP and with four cytosolic enzymes under identical settings (2,105 pairs). iLIS ≥ 0.223 called 3.3% of these controls positive (95% CI 2.6 to 4.2%; **Figure S16**), so truncation does not raise the false positive rate above that of full-length controls. On the same basis (AF rank-1 model, isoform pairs), 9.3% of the screen was called positive (11.8% at the gene level with the highest-iLIS model; **Table S8**). The ECD controls come from a different cellular compartment and do not represent an extracellular background, so the difference between 9.3% and 3.3% is not a measure of enrichment.

For pairs of extracellular proteins, no verified negative set exists, because secreted proteins and receptor ectodomains can meet outside the cell. Pairs presumed to be non-cognate (ECDs paired with Wg, or with ligands whose receptor is known) were called at 11%, compared with 9.6% for the Wg controls of the calibration set and 9.3% for the screen overall. Some of these pairs may be uncharacterized interactions, so the false positive rate among extracellular pairs is not established, and the screen ranks candidate receptors for a given ligand.

Among the heterodimeric PPIs, 15,386 are supported by fly literature evidence, 28,268 by interolog conservation (DRSC Integrative Ortholog Prediction Tool [DIOPT] score ≥ 2; Hu et al., 2011), and the remaining 101,125 (70% of positive heterodimers) have neither a curated fly-interaction record nor conserved-interolog support at DIOPT ≥ 2, representing candidate interactions without prior physical-interaction evidence (**Table S9**, evidence class “unreported”), with these evidence classes broken down by prediction source in **Table S10**. Most unreported positives come from the ligand-receptor and kinase-related targeted screens.

None of the validation analyses in this study tests this class directly: the benchmark, recovery, and network analyses exclude it by construction, and it enters the allele analysis only through pooled residue frequencies. In independent annotation, unreported positives shared a specific Gene Ontology (GO) cellular-component term more often than sub-threshold unreported pairs (96 of the 85,845 positives evaluable for this annotation; rate ratio 1.39, 95% CI 1.12 to 1.74; odds ratio 1.32, 1.05 to 1.66, after stratification by hub protein) but not shared protein-complex terms, coexpression, or text-mining associations, and interaction records added after our evidence snapshot were less frequent among them (6 pairs; rate ratio 0.43, 95% CI 0.19 to 0.99; **Table S4**). With base rates near 0.1%, none of these comparisons constrains the precision of the class in either direction. The fraction of unreported candidate pairs called positive (12.2%; 101,190 of 832,750) also lies within the range of control false positive rates (3% to 20%), so the call rate alone gives no evidence of enrichment in this class. The precision of this class is therefore unresolved, and these pairs are confidence-ranked candidates, not established interactions.

To explore the functional organization of the positive PPIs, we selected those supported by the fly literature or by conserved interologs. Because interolog inference can be permissive at low ortholog confidence, we applied a more stringent threshold of DIOPT ≥ 4 (rather than the ≥ 2 threshold used for database annotation) for network construction, reducing the 28,268 interolog-supported pairs to 8,265. Retaining proteins with at least two interactions then yielded a network of 5,224 proteins and 21,657 PPIs (**Figure 4B**). Leiden community detection (Traag et al., 2019) identified 31 clusters (**Figure 4C**), and GO enrichment analysis showed that most clusters have a characteristic top-ranked biological process (30 different top terms across the 31 clusters), although each cluster retains several enriched terms and a few terms are shared between clusters (**Figure S17**; **Table S11**).

To resolve functional heterogeneity within each cluster, we performed recursive sub-clustering, yielding 257 sub-clusters and 407 sub-sub-clusters (see Methods; **Figure S18**). We highlight four examples below that illustrate how this hierarchical analysis resolves pathway-level modules into individual protein complexes. Full results are available in **Tables S11-S15** and the interactive network viewer (https://flyark.github.io/FlyPredictome-network).

In Cluster 1 (kinases and transcriptional regulators), sub-cluster C1-4 (JNK cascade and growth control) contains a 14-protein sub-group (C1-4-2) corresponding to the core Hippo pathway components (the kinases Hpo and Wts, the Wts co-activator Mats, and the scaffold protein Sav) together with upstream regulators Ex and Jub, the Sd co-repressor Tgi, and the MAP4K family kinase Hppy (**Figures 4D**, **S19**). This sub-group also includes CG12645, a predicted zinc-binding protein localized to late endosomal and lysosomal membranes, which is predicted to bind both Sav (iLIS = 0.535) and the upstream regulator Kibra (iLIS = 0.342), suggesting a potential role in membrane-associated Hippo pathway regulation.

In Cluster 10 (signal transduction and cytoskeleton), sub-cluster C10-1 (Arp2/3-mediated actin nucleation) contains a 15-protein sub-group (C10-1-3) comprising two structurally linked sub-modules: Arp2/3 complex subunits (Arp2, Arpc1, Arpc4, Arpc5) with the WASp actin regulatory module (WASp, Vrp1, Nwk), and Eph-Ephrin signaling components (Ephrin, Dock) (**Figures 4E**, **S20**). The edges linking these two sub-modules are supported by interolog conservation, consistent with the established role of Eph receptor signaling in driving actin remodeling through N-WASP and Cdc42 (Irie and Yamaguchi, 2002).

In Cluster 12 (autophagy), the macroautophagy sub-cluster C12-1 separates the PI3K complex required for phagophore nucleation (Atg6, Vps15, Pi3K59F, Atg14 in C12-1-3) from a 10-protein module (C12-1-2) (**Figures 4F**, **S21**). The latter contains the Atg1 kinase complex (Atg1, Atg13, Atg17/FIP200), the Atg8-processing proteases Atg4a and Atg4b and their substrate Atg8b, the PI3P-binding proteins Atg18a and Atg18b, the lipid transfer protein Atg2, and the selective autophagy receptor ref(2)P (p62/SQSTM1 ortholog).

In Cluster 14 (chromatin and DNA repair), sub-cluster C14-4 (DNA double-strand break repair) contains a 10-protein sub-group (C14-4-2) comprising the non-homologous end joining (NHEJ) factors Ku heterodimer (Irbp and Ku80) and DNA Ligase IV, together with Parp, the RPA complex (RpA-70, RPA2, RPA3), and Blm helicase (**Figures 4G**, **S22**; **Figure 4D-G** is enlarged in **Figure S23**). This module also includes CG3448, a distant ortholog of human XRCC4, predicted to bind DNAlig4 (iLIS = 0.448). XRCC4 is the obligate partner of DNA Ligase IV in the NHEJ ligation complex across eukaryotes (Sibanda et al., 2001). CG3448 has not been functionally characterized in *Drosophila*, but its co-purification with DNAlig4 in AP-MS (Bhat et al., 2024) and the structural prediction here both support its assignment as the fly XRCC4.

### Higher-order protein complexes from network sub-clusters

Having partitioned the interactome into functional modules by their pairwise interactions, we next asked whether these modules assemble into single higher-order complexes. We first folded three representative sub-clusters with AlphaFold3 (AF3; Abramson et al., 2024) and, where an experimental structure was available, compared the predictions to it. Sub-cluster C6-13 comprises Orc1 to Orc5 of the origin recognition complex (ORC) together with Cdc6. Folding these subunits alone with AF3 (without the origin DNA present in the experimental structure) recovered an assembly closely matching the cryo-EM structure of *Drosophila* ORC (PDB: 7JGR) at 1.43 Å Cα root-mean-square deviation (RMSD; TM-score 0.959), with per-subunit TM-scores of 0.957 to 0.997 (**Figure 5A,B**), showing that the quaternary arrangement, not only the individual folds, was recovered from the sub-cluster membership alone (this structure was deposited before the AlphaFold3 training cutoff).

**Figure 5.**
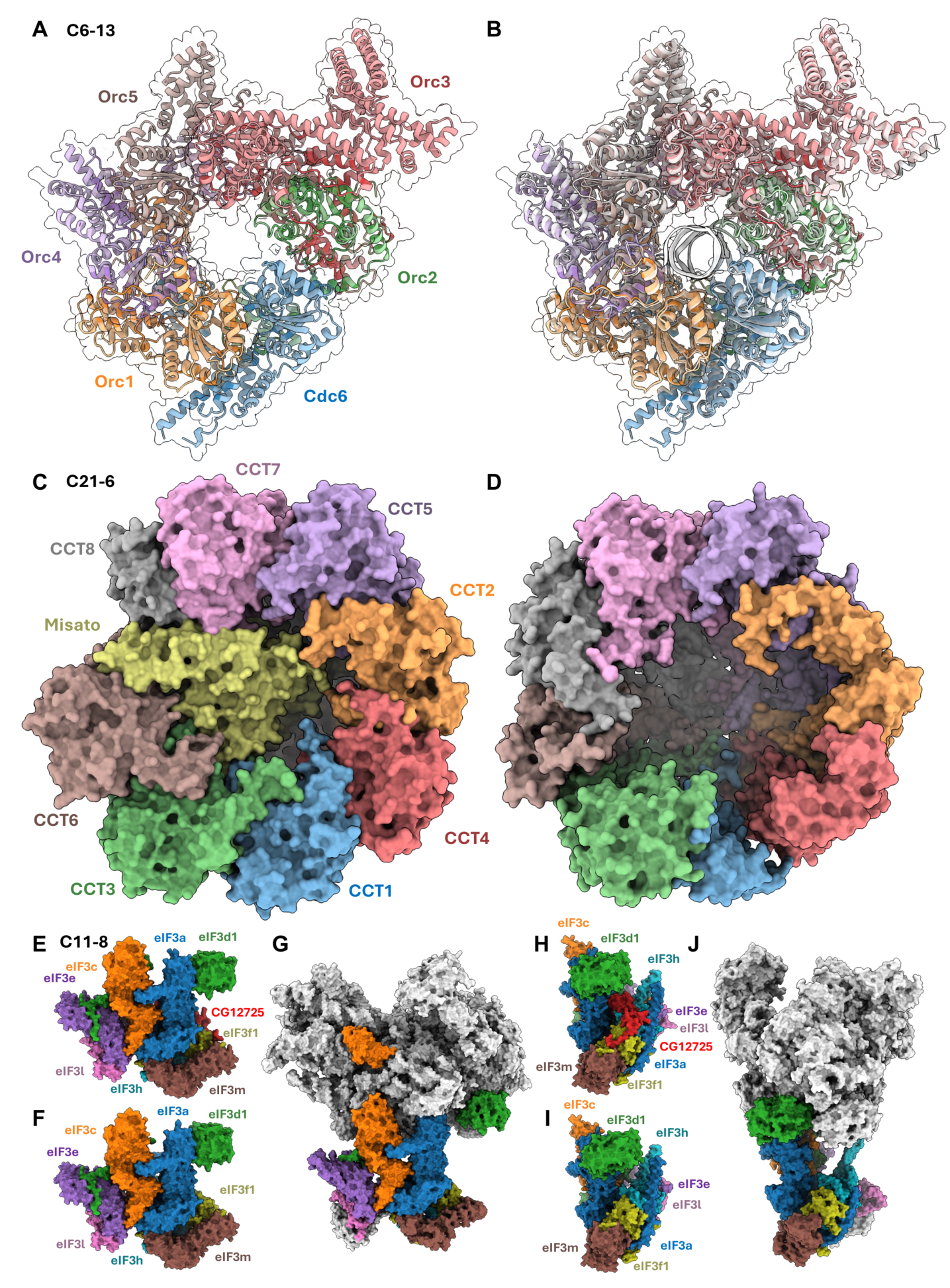
Network sub-clusters predict higher-order protein complexes. **(A)** AlphaFold3 prediction of the ORC-Cdc6 assembly (sub-cluster C6-13: Orc1 to Orc5 and Cdc6), colored by subunit. Detailed subunit-pair interactions are shown in **Figure S24A**. **(B)** The same prediction superimposed on the *Drosophila* ORC cryo-EM structure (gray; PDB: 7JGR), which resolves ORC bound to origin DNA and Cdc6; predicted without the DNA, it recovers the assembly (whole-complex Cα RMSD 1.43 Å, TM-score 0.959; per-subunit TM-scores 0.957 to 0.997). **(C)** AlphaFold3 prediction of the CCT/TRiC chaperonin with Misato (sub-cluster C21-6: CCT1 to 8 with Misato), viewed down the ring axis; the closed CCT1 to 8 ring matches one ring of the human TRiC cryo-EM structure (PDB: 7LUM), with Misato lodged in the central folding cavity, contacting most of the eight CCT subunits (most extensively CCT6). Detailed subunit-pair interactions are shown in **Figure S24A**. **(D)** AlphaFold3 prediction of the eight CCT subunits folded alone, without Misato, viewed down the same ring axis; the ring closes into the same architecture, and the central cavity that Misato occupies in **C** is left open. **(E, F)** AlphaFold3 prediction of the nine-subunit eIF3 assembly (sub-cluster C11-8), colored by subunit, shown with (**E**) and without (**F**) the uncharacterized ubiquitin-like protein CG12725 (red subunit). CG12725 co-purifies with eIF3 subunits in the DPiM2 AP-MS dataset (Bhat et al., 2024) and is absent from all deposited eIF3 structures; the prediction positions it against eIF3a, eIF3f1, and eIF3h. Detailed subunit-pair interactions are shown in **Figure S24A**. **(G)** The human 48S pre-initiation complex cryo-EM structure (PDB: 28ZX, extended RCSB accession pdb_000028zx), with its eIF3 subunits colored as the corresponding subunits in **E**/**F** and all other components in gray; eight of the nine predicted subunits have a counterpart in the reference, and CG12725 has none. **(H, I)** A second orientation of the eIF3 assembly, shown with (**H**) and without (**I**) CG12725. **(J)** The reference structure of **G** in the second orientation.

Because ORC’s contact map is experimentally known, it also shows how pairwise metrics behave when they are read from a multi-subunit model. Within the AF3 assembly, three non-contacting subunit pairs (Cdc6-Orc5, Orc1-Orc3, and Orc2-Orc4) score zero by iLIS yet reach 0.53 to 0.59 by ipSAE and 0.68 to 0.74 by ipTM (**Figure S24A**; values in **Table S4**), because a multimer model positions all subunits confidently relative to one another whether or not they touch. When the same pairs were predicted individually with AFM, iLIS and ipSAE both scored them near zero (**Figure S24C**). Across 1,226 subunit pairs predicted both ways, the four metrics behaved alike in pairwise predictions and diverged only in the co-folded assemblies, where non-contacting pairs passed the iLIS and actifpTM* cutoffs about three times less often than the ipSAE and ipTM cutoffs (17% and 18% versus 57% and 49%; **Figure S24B-E**). The pairwise models are AlphaFold-Multimer predictions and the assemblies are AlphaFold3 predictions, so the two arms differ in predictor as well as in context, whereas the metrics within the assemblies are compared on the same models. The divergence among the metrics within the assemblies therefore reflects whether a metric incorporates an intermolecular distance filter (Cβ-Cβ ≤ 8 Å in iLIS and actifpTM*, absent in ipSAE and ipTM) when scoring subunit pairs inside multimers, rather than performance on dimers.

Beyond reproducing complexes with known structures, the same approach can model an engagement that no experimental structure has resolved. Sub-cluster C21-6 groups the eight-subunit CCT/TRiC chaperonin (CCT1 to 8) with Misato (mst), which co-purifies stoichiometrically with all eight subunits and was proposed, from comparative modeling of its tubulin-like fold, to occupy the chaperonin’s folding cavity (Palumbo et al., 2015). Co-folding the nine proteins assembles the closed CCT ring with Misato lodged in that cavity, in contact with most subunits and most extensively with CCT6 (**Figure 5C**, **S24A**). The prediction places all nine chains simultaneously, converting a decade-old hypothesis into a specific, testable interface. Folded without Misato, the eight CCT subunits closed into the same ring and left the central cavity open (**Figure 5D**), so ring closure in the model does not require Misato.

AlphaFold3 prediction can also place a potential accessory subunit. Sub-cluster C11-8 contains the eIF3 translation-initiation complex together with CG12725, an uncharacterized ubiquitin-like protein reported as an eIF3 interactor by AP-MS (Bhat et al., 2024) but absent from every deposited eIF3 structure. Folding this eIF3-CG12725 core (nine subunits) positions CG12725 against the eIF3a, eIF3f1, and eIF3h subunits, supplying a possible structural location for an interaction previously known only as a co-purification edge and raising the possibility that this ubiquitin-like protein transiently associates with the eIF3 core (**Figure 5E-J**, **S24A**). Subunit placements absent from every deposited structure (Misato in the CCT folding cavity, CG12725 within eIF3) should be interpreted with caution as structural hypotheses until tested directly; in both cases, the association itself was previously observed (Palumbo et al., 2015; DPiM2 dataset of Bhat et al., 2024), and what the prediction adds is an explicit interface.

More broadly, we extended this approach to 398 assemblies across the network sub-clusters, folding their subunits together rather than pairwise. Of the 202 with an evaluable cryo-EM reference (**Figure S25A**), 178 (88%) recovered every subunit-subunit contact observed in the deposited structure (median recall 1.00, mean 0.98; contacts defined as Cβ-Cβ ≤ 8 Å, scored over the subunits shared between prediction and reference). The rate was the same for references released after the AlphaFold3 training cutoff of 2021-09-30 (124 of 141) and before it (54 of 61). This rate is conditional on an assembly having folded into a connected complex and on having an evaluable reference: the 398 assemblies were retained from 2,859 folded AlphaFold3 jobs (the 178 fully recovered assemblies are 45% of the former and 6.2% of the latter), most are subsets or variants of their sub-cluster, and at the level of network modules, 76 of the 257 sub-clusters (30%) have at least one assembly that recovered every deposited contact (selection rules in Methods and **Figure S25A**). Complete recovery declined monotonically with assembly complexity, from 98% for three-subunit assemblies to 75% for those with eleven or more subunits (**Figure S25B**), and from 97% for assemblies with one to three deposited contacts to 56% for those with more than twenty (**Figure S25C**).

Neither nucleic acid nor ligands in the deposited reference impaired recovery; references containing ligands in fact recovered better than those without (144 of 158, 91%, versus 34 of 44, 77%, two-sided Fisher’s exact P = 0.02; nucleic acid, P = 0.26), so the basis of incomplete recovery, beyond assembly size, remains unresolved. A full catalog of folded assemblies, spanning both structurally supported and previously uncharacterized complexes, is provided in **Table S16**; the same assemblies, with per-subunit interface-confidence scores and 3D coordinates, are available in the interactive network viewer through LIVIA (Local Interaction Visualization and Analysis) (Kim and Perrimon, 2026).

### Interaction fingerprints resolve partner-specific surfaces

Because FlyPredictome maps each prediction to the residues at its interface, the contact residues a protein uses across all of its predicted partners can be assembled into a clustered interaction fingerprint that decomposes its surface into partner-specific binding regions, an analysis that conventional node-and-edge interactomes cannot support. As an example, the interactive FlyPredictome Network Explorer (**Figure S26A**) presents the predicted partners of the transcriptional cyclin-dependent kinase Cdk12 as a node-and-edge ego-network (**Figure S26B**). The clustered interaction fingerprint resolves these partners to residue level, where they cluster by shared contact residues: the cluster containing the transcriptional cyclins CycK, CycT, and CycH converges on the C-terminal protein-kinase domain (residues 804 to 1098, a folded, high-pLDDT region), whereas other clusters engage separate N-terminal segments predicted with low confidence (**Figure S26C**). Projected onto the Cdk12 structure these clusters occupy spatially distinct surfaces (**Figure S26D**). This decomposition tracks each protein’s architecture rather than being specific to Cdk12: for proteins as varied as the transcription factor Myc, the scaffold Armadillo/β-catenin, the receptor Egfr, and the Notch DNA-binding factor Su(H), partners resolve onto distinct folded domains, a repeat groove, or several discrete surfaces (**Figure S27**). Such partner-resolved fingerprints are available for 5,004 of the 5,224 proteins in the evidence-supported network.

## DISCUSSION

FlyPredictome provides a proteome-scale structural interactome for *Drosophila melanogaster*.

By scoring 1.5 million pairwise AFM predictions with iLIS, we identified positive PPIs across experimentally reported and computationally inferred interaction categories, providing binding interfaces that predict which domains and residues mediate each interaction. Furthermore, by scoring only the confident interaction domain within the full-length proteins, iLIS extends this capability to interactions involving proteins with extensive low-confidence regions, which are difficult to detect with global confidence metrics. Many such regions are likely intrinsically disordered, a class of sequence that mediates many key signaling events in eukaryotes (van der Lee et al., 2014; Wright and Dyson, 2015). Because iLIS requires intermolecular contact in the model, each positive prediction specifies a contact interface, and combining these calls with prior interaction evidence yielded an evidence-supported PPI network.

The residue-level detail of these predictions enabled us to demonstrate that phenotype-associated missense mutations are enriched at predicted interaction sites, supporting the functional relevance of these structural models. The enrichment at direct-contact residues persists after adjustment for burial, conservation, prediction confidence, and MSA depth, whereas the graded enrichment across the broader interaction domain largely tracks prediction confidence (**Figure S12**). This synergy between genetics and structural prediction is exemplified by the Fz-Dsh interface, where classical fly genetics had identified PCP-specific DEP-domain mutations whose structural consequences are now visualized at residue resolution by AFM predictions.

The resource has also been tested prospectively. In FlyPredictome, Beat-Ia and Side (iLIS = 0.511, predicted in June 2023; nine further Beat-Side paralog pairs are positive in **Table S9**) interact through their N-terminal immunoglobulin (IG1-IG1) domains, matching the IG1-IG1 binding topology subsequently resolved in the crystal structure of the paralogous Beat-Vc-Side-VI complex (PDB: 9NSF), with structure-guided interface mutations showing that the Beat-Ia-Side interaction is required for motor neuron-muscle wiring *in vivo* (Priest et al., 2026). The predicted Fz-Dsh complex likewise matched a cryo-EM structure published after the AFM training cutoff (**Figure S13A**).

Grouping proteins by their evidence-supported predicted interactions recovers functional modules; folding a subset of these with AlphaFold3 reconstructs known assemblies and proposes structures for complexes that no experiment has resolved, indicating that the modules the binary predictions define can contain genuine higher-order assemblies. In the opposite direction, because every interaction is mapped to its interface residues, the partners of a single protein can be resolved onto distinct binding surfaces, distinguishing interactors that share a predicted surface from those that engage separate regions.

Several limitations of the current FlyPredictome dataset should be noted. Coverage is not exhaustive: the 1.5 million predictions were assembled from targeted prediction sets rather than from an all-by-all screen of the *Drosophila* proteome, so the large majority of possible pairs have not been tested and interactions between proteins not previously linked by any evidence remain largely unexplored. This bias is compounded in the network layer, which is built from interactions supported by existing fly literature or conserved interologs and therefore underrepresents pathways for which experimental PPI data are sparse; assignments based on few edges should be interpreted accordingly.

Our AFM predictions also do not account for post-translational modifications, cofactors, or cellular compartmentalization, and are limited to binary interactions. As a result, interactions that depend on these factors, or that form only within a larger assembly, may be missed by pairwise prediction, even though folding a module’s proteins together can recover the assembly itself (**Figure 5**); this contributes to the modest recovery rates observed for experimental PPI datasets. Conversely, structurally compatible protein pairs that do not co-localize or co-express *in vivo* should be interpreted with caution.

The benchmark also indicates why metrics disagree on individual pairs and where iLIS is limited. Pairs called by iLIS but not by ipTM or pDockQ typically combine long, low-confidence chains with a compact interface, as expected for short motifs docking onto folded domains. In such cases, surrounding sequence dilutes global and all-contact averages, unlike domain-restricted scoring. However, controls with this architecture are also called more often. Conversely, iLIS misses interfaces with uniformly moderate confidence (inter-chain PAE near or above 12 Å) and interactions lacking direct model contacts (**Figure S6M-P**; **Table S1**), and at the 1% FPR cutoff on single models, actifpTM*-only calls were more often correct than iLIS-only calls (**Table S4**).

The threshold for positive PPI classification may need adjustment for different biological contexts, and its false positive rate depends on the negative population: iLIS ≥ 0.223 passed between 3% and 20% of controls across the control classes examined here (**Figures S4B, S16**; **Table S6**). For two parts of the atlas, precision remains unestimated. The precision of the unreported positives, which make up 70% of positive heterodimers, is unresolved, and these pairs are confidence-ranked candidates. In the ligand-receptor screen, extracellular-domain controls were called at 3.3%, but no verified negative set exists for pairs of extracellular proteins, so the false positive rate within the screen is not established and its calls rank candidate receptors for a given ligand.

No single confidence metric is uniformly best: agreement among the four metrics was higher on the benchmark than across the atlas, where many predictions are borderline (**Figure S7A, C**). Rather than relying on a single choice, FlyPredictome also provides a composite tier score that aggregates five metrics; it recovers the calls shared by the individual metrics without being driven by any one of them (**Figure S7B, D**), and it is available on the database (Methods). On held-out pairs, the composite score performed like iLIS and did not exceed it (**Table S4**). Because every analysis in this study starts from iLIS calls, pairs below the iLIS cutoff that only other metrics support are absent from the positive set and from the network. For an individual pair of interest, the database reports the full set of metrics and the composite score in addition to the binary call.

FlyPredictome provides all precomputed predictions as an open resource, including those below the positive threshold, rather than reporting only high-confidence interactions. These scores help narrow down candidate interaction partners, while residue-level interface annotations can inform the design of interaction-specific mutations or domain truncations that selectively disrupt individual binding events. The database also provides specialized prediction sets, including kinase interactome and ligand-receptor prediction sets, to serve as a resource for signaling and cell communication studies, and will continue to expand as new prediction sets are added.

Because it scores structural confidence alone, iLIS can also be applied to settings with no prior interaction data, including higher-order assemblies of three or more subunits and *de novo* designed binders. While computational binder screens are typically ranked by confidence alone, ranking natural complexes by an interface area-weighted metric such as iLISA prioritizes established partners with extensive interfaces (**Figure S10**); incorporating this interface-extent axis could similarly help prioritize *de novo* designs that bury substantial surface area. The scoring, validation, and allele enrichment framework described here can likewise be applied to any organism with proteome-scale structural predictions and a catalog of phenotype-associated variants.

## METHODS

### AlphaFold-Multimer predictions

Protein structure predictions were performed using LocalColabFold (ColabFold v1.5.2 to v1.5.5; Mirdita et al., 2022) with the AlphaFold-Multimer v2.3 model weights (ColabFold model_type alphafold2_multimer_v3, distributed with AlphaFold v2.3; Evans et al., 2021; Google DeepMind, 2022). All pairwise predictions in this study, including every benchmark and control set, used these same v2.3 weights; no predictions were made with the v2.1 (multimer_v1) or v2.2 (multimer_v2) weights, which differ in their handling of chain overlaps and clashes. Templates were not used, multiple sequence alignments (MSAs) were built in unpaired_paired mode, and no structure relaxation was applied. MSAs were generated with MMseqs2, either through the ColabFold MSA server (early runs with ColabFold v1.5.2) or with colabfold_search against UniRef30 (version 2202) and the ColabFold environmental database (later runs). All computations were performed on the Harvard O2 high-performance computing cluster and the Longwood high-performance compute cluster using a variety of NVIDIA GPUs (TeslaV100, RTX6000, RTX8000, A40, A100, A6000, L40S, and H100). Each prediction generated five models with five recycling iterations. Interaction confidence was assessed using the integrated Local Interaction Score (iLIS).

### Integrated Local Interaction Score

iLIS is defined as the geometric mean of LIS and cLIS: iLIS = √(LIS × cLIS). LIS (Local Interaction Score) quantifies the confidence of the predicted interaction domain by inversely rescaling inter-chain PAE values within the confident region (PAE ≤ 12 Å) to a 0-to-1 score, where PAE = 12 Å maps to 0 and PAE = 0 Å maps to 1. LIS is computed as the mean of these rescaled values over all inter-chain residue pairs with PAE ≤ 12 Å, averaged over both chain orientations of the PAE matrix. Throughout, the c-prefix (cLIS, cLIA, cLIR) denotes the corresponding quantity restricted to residue pairs in direct intermolecular contact (Cβ-Cβ ≤ 8 Å) within the confident interaction domain (PAE ≤ 12 Å); cLIS is therefore the mean of the same rescaled values over that contact subset.

This contact filter is needed because in some AFM predictions two protein chains can be positioned in proximity in 3D space without forming genuine intermolecular contacts, resulting in spuriously high LIS values. The geometric mean formulation balances domain-level and contact-level confidence, penalizing predictions where LIS is high but no direct contacts exist (cLIS = 0; **Figure S1B**). This formula was defined a priori because additive combinations (such as LIS + cLIS) remain high when LIS is large even if cLIS = 0, whereas √(LIS × cLIS) collapses to zero whenever direct contacts are absent.

Across alternative formulations, the arithmetic mean, harmonic mean, minimum, and LIS gated by cLIS > 0 performed within 1 to 2 percentage points of the geometric mean at 10% and 5% FPR (AUC differences ≤ 0.003), whereas LIS alone recovered 6.7 points fewer interactions at 1% FPR and cLIS alone 3.1 points fewer at 5% FPR (paired bootstrap confidence intervals excluding zero; **Figure S2A-D**; **Table S4**, sheet G), showing that using both components matters more than the specific functional form.

Previously, this issue was addressed using a double-cutoff approach requiring both LIS and LIA (Local Interaction Area) thresholds (Kim et al., 2024), an approach that has since been independently adopted (Qi et al., 2026). iLIS simplifies this to a single integrated score. Contact-filtered cLIS and the geometric-mean iLIS were released as open-source code in the AFM-LIS repository (cLIS in May 2024 and iLIS in March 2025; https://github.com/flyark/AFM-LIS) and are formally defined, calibrated, and benchmarked in the present study. By the same design, iLIS scores only direct contacts in the model: an interaction that forms no direct residue-residue contact between the two chains, for example a genuine association bridged through a shared partner or a subunit correctly positioned within an assembly but not touching a given neighbor, has cLIS = 0 and therefore scores zero.

The PAE ≤ 12 Å window (together with the earlier LIS and LIA cutoffs) was originally selected on the fly and human Y2H datasets and tested on the yeast Y2H and ELM datasets in Kim et al. (2024). Here, all four datasets were pooled to compare metrics, and the yeast, fly, and human Y2H sets were used to calibrate the proteome-scale threshold (holding out the ELM and shuffled sets; see below), so the full-length benchmark overlaps that earlier tuning whereas the yeast and ELM subsets and the post-cutoff PDB benchmark (**Figure S5**) were not used to select the 12 Å window.

Because this threshold is applied as a hard window, and inter-chain pairs are rescaled so that a pair at PAE = 12 Å contributes zero, iLIS is insensitive to interfaces that are extensive but predicted with uniformly low confidence: an interface whose inter-chain PAE values all lie just outside the window yields no qualifying residue pairs and therefore a score of zero, irrespective of its area. Interactions of this kind are systematically excluded rather than merely down-weighted. Users interested in weak or low-confidence engagements can recompute the score with a more permissive window (for example PAE ≤ 15 Å) at the cost of specificity; the threshold is exposed as a parameter in the iLIS analysis pipeline (https://github.com/flyark/AFM-LIS).

### Other confidence metrics

In addition to iLIS, we benchmarked multiple confidence metrics. Local confidence metrics restrict scoring to the high-confidence interaction region: iLIS, LIS (Kim et al., 2024), cLIS, actifpTM (Varga et al., 2025; denoted actifpTM* in our post-hoc implementation), and ipSAE (Dunbrack, 2025). Global confidence metrics score all inter-chain residue pairs regardless of confidence: ipTM, Model Confidence (Evans et al., 2021), and ifPAE (the mean inter-chain PAE across all residue pairs). Contact-based metrics score residue pairs in direct intermolecular contact, selected by distance rather than by a PAE domain window: pDockQ (Bryant et al., 2022), pDockQ2 (Zhu et al., 2023, which additionally incorporates PAE as a value in its score), and ifPAE_d8 (mean inter-chain PAE restricted to Cβ-Cβ ≤ 8 Å pairs).

Within the local group, the metrics differ further in whether a distance criterion is applied. iLIS incorporates a direct-contact requirement by taking the geometric mean of LIS (inter-chain PAE ≤ 12 Å) and contact-filtered cLIS (PAE ≤ 12 Å and Cβ-Cβ ≤ 8 Å). actifpTM weights inter-chain TM-score contributions by contact proximity (Varga et al., 2025).

Because the original actifpTM is computed inside the AlphaFold2 confidence module from predicted contact probabilities (Cβ-Cβ < 8 Å) and aligned-error bin probabilities, which are not saved by default in ColabFold, AlphaFold3, or AlphaFold Database outputs, our pipeline (lis.py) computes a post-hoc approximation denoted actifpTM* directly from the output structure and PAE matrix: for each residue forming at least one inter-chain Cβ-Cβ < 8 Å contact (Cα for glycine), it averages 1 / [1 + (PAE / d_0_)^2^] over its contacting partner residues using the full-complex scaling length d_0_ = 1.24(max(L_total, 19) − 15)^(1/3) − 1.8 (zero if no inter-chain contacts exist), and takes the maximum across contacting residues in both chain directions. ipSAE selects residue pairs by inter-chain PAE alone (Dunbrack, 2025; computed throughout with the d0res definition, PAE < 10 Å, taking the larger of the two chain directions), and ipTM applies no selection at all, so neither requires the residues it scores to be in contact.

For pairwise-versus-multimer comparisons (**Figure S24**), 1,226 cryo-EM-referenced subunit gene pairs were matched by FlyBase gene ID (FBgn) to atlas AF rank-1 models (97% of chains sequence-identical): metrics incorporating a distance criterion (iLIS and actifpTM*) report modeled contact in both dimers and multimers, whereas positional metrics (ipSAE and ipTM) report direct interaction in dimers and co-complex placement in multimers.

Two area-weighted variants report interaction area rather than confidence alone: iLIA = √(LIA × cLIA), the geometric mean of the Local Interaction Area (LIA, the number of inter-chain residue pairs with PAE ≤ 12 Å) and the contact LIA (cLIA); and iLISA = iLIS × iLIA, which scales interaction confidence by interaction area. We also evaluated composite tier scores across five metrics (iLIS, iLISA, actifpTM*, ipSAE, ipTM): each metric contributes one, two, or three points as a pair passes its deployed 10%, 5%, or 1% FPR threshold (**Table S6**, sheet Deployed_cutoffs; for iLISA, the deployed tiers differ from the complete-case recalibration listed in the FPR_thresholds sheet), yielding composite15 (0 to 15) on the single best model (**Figure S7B, D**) and composite30 (0 to 30) when summed over the single-model value and the five-model average, reported alongside the individual metrics in the database (**Table S9**).

For ifPAE and ifPAE_d8, lower values indicate higher-confidence interfaces; inverted values were used in correlation and ROC analyses for consistent directionality. All benchmark metrics were evaluated using both the AF rank-1 model (AlphaFold’s own ranking by 0.8 × ipTM + 0.2 × pTM) and the average across all five models per pair. Spearman correlation analysis for the benchmark dataset is available in **Figure S3A-B**. All benchmarked metrics, with their scoring basis, PAE and contact filtering, and benchmark performance, are compared in **Table S2**. Bootstrap confidence intervals and FPR thresholds are provided in **Tables S3** and **S6**, respectively.

### Benchmark dataset construction

The benchmark dataset comprised a positive reference set and multiple control sets. The positive set (736 PPIs) integrates four sources: the human Positive Reference Set (PRS; Braun et al., 2009), the fly PRS from the FlyBi Y2H screen (Tang et al., 2023), the yeast PRS (Yu et al., 2008), and structure-validated interactions from the Eukaryotic Linear Motif (ELM) database (Kumar et al., 2024).

The control set (5,909 PPIs) includes Random Reference Sets (RRS) from the three Y2H screens (protein pairs not reported to have an interaction at the time of curation), GFP controls (each PRS protein paired with GFP, serving as a non-interacting folded protein), subcellular compartment controls (PRS proteins paired with the secreted protein Wingless, testing whether compartment mismatch alone produces low scores), and separately shuffled pairs from each PRS and ELM set (pairing proteins with non-cognate partners).

The complete benchmark dataset (6,645 source-level records: 736 positives and 5,909 controls, corresponding, after removal of reciprocal and cross-source duplicates and of nine control records (seven self-pairs and two shuffled pairs that are also reference positives), to 713 unique positives [383 Y2H PRS and 330 ELM] and 5,890 unique controls [1,270 in RRS, PRS + GFP, and PRS + Wg, of which 1,255 carry complete multi-metric scores for calibration, and 4,620 in shuffled and ELM held-out classes, totaling 4,635 non-calibration controls]) is provided in **Table S1**.

A time-split validation set of two-chain complexes (mostly heterodimers, with a minority of homodimers) whose experimental structures were deposited in the PDB after the AFM training cutoff (2021-09-30) was predicted from the deposited-construct (SEQRES) sequences and, for the 198 predictions included in the DockQ analysis, from full-length sequences with AlphaFold-Multimer (five models per prediction). Every model was scored by DockQ (v2; Mirabello and Wallner, 2024) against its deposited structure (chains mapped by sequence), and the iLIS of the highest-iLIS model was correlated (Spearman ρ and Pearson r) with DockQ both with one prediction per PDB entry (SEQRES preferred; 402 complexes) and over all 536 predictions (**Figure S3K**).

To test whether the correlation depends on similarity to the training set, each query chain was aligned by BLASTp to all protein chains of the 182,459 PDB entries released on or before the cutoff (618,237 chains); complexes were stratified by the maximum sequence identity of either chain to any such chain, and the correlation was recomputed within each tier; it persisted for the 49 complexes with < 30% identity to any training-era chain (ρ = 0.70; 65 predictions, ρ = 0.62). Sequence redundancy among the complexes was additionally controlled by BLASTp clustering (≥ 30% identity on both chains) and recomputing on one representative per cluster (ρ = 0.75 over predictions).

Per-pair iLIS and DockQ values for all time-split complexes, with PDB deposition era and redundancy-cluster assignment, are provided in **Table S5**. For the DockQ analysis, the model with the highest iLIS among the five was used, as for atlas calls (see Candidate pair selection); for the classification analysis below, the AF rank-1 model (ranked by AlphaFold’s model confidence, 0.8 × ipTM + 0.2 × pTM) or the five-model average was used, identically for every metric.

For the independent classification benchmark (**Figure S5**), positives were the 338 after-cutoff complexes predicted from SEQRES sequences and negatives were 1,257 controls pairing the proteins of these complexes with GFP (629) or Wg (628), predicted with the same settings. ROC-AUC, precision-recall AUC (average precision), and sensitivity at matched 10%, 5%, and 1% FPR were computed for iLIS, LIS, cLIS, iLISA, actifpTM*, ipSAE, ipTM, pDockQ, and pDockQ2, with 95% confidence intervals and paired differences from 2,000 stratified bootstrap replicates with shared resampling indices.

Homology to the training era was assessed by BLASTp of each chain against all PDB chains released on or before 2021-09-30 (e-value ≤ 10^-3^, alignment covering ≥ 50% of the shorter sequence), taking the higher of the two chains’ maximum identities; complexes were stratified at < 30%, 30 to 90%, and ≥ 90% identity, and complex-level relatedness was assessed by whether a single training-era entry contained homologs of both partners (229 of 338); similarity at the level of the interface was not assessed. Redundancy was reduced by keeping one randomly chosen complex per sequence cluster (fixed seed). The deployed cutoffs were applied without recalibration to report achieved sensitivity and FPR with Wilson 95% confidence intervals, overall, by control type, and by pLDDT subgroup. The 140 full-length re-predictions (134 of which could be scored by DockQ and are included in **Figure S3K**; for the other six, sequence differences between the full-length sequence and the deposited construct prevented the automatic chain mapping) were analyzed in the same way against 313 full-length GFP/Wg controls as a secondary set, on which iLIS and ipTM had the same ROC-AUC (0.841; **Figure S5D, E**). The controls of this benchmark pair each protein with GFP or Wg and do not include non-cognate pairs of related proteins.

### Threshold calibration for positive PPI classification

Thresholds for positive PPI classification were calibrated using ROC analysis on the Y2H-derived PRS from yeast, fly, and human screens as positives, and three control sets as negatives: RRS, PRS proteins paired with GFP, and PRS proteins paired with Wingless (Wg, a secreted protein serving as a subcellular compartment control). ELM interactions and randomly shuffled pairs were not used for calibration; they were held out and used to test the fixed cutoffs (see below). Including them among the calibration negatives raises the 10% FPR cutoff and leaves the 1% FPR cutoff unchanged, because motif-hub and shuffled pairs exceed a given iLIS more often than RRS, GFP, and Wg pairs (**Figure S4B**): the 10% FPR cutoff of iLIS rises from 0.223 to 0.288, sensitivity on Y2H PRS pairs falls by about 2 points, and 23.8% of the fly-literature positives and 32.4% of the interolog-supported positives in the atlas would no longer be called (**Table S4**, sheet D). For proteome-scale candidate discovery, we retained iLIS ≥ 0.223 (10% FPR) alongside the 5% (≥ 0.339) and 1% (≥ 0.551) tiers and five-model averages. Pairs lacking a score for any benchmarked metric were excluded so that all metrics were calibrated on an identical set, leaving 357 positives and 1,255 negatives.

Thresholds were set by target false positive rate (FPR), the fraction of calibration controls (RRS, GFP, and Wg) exceeding the cutoff (**Table S6**). FPR depends on the metric, the threshold, and the negative population used to estimate it, but, unlike the false discovery rate (FDR), not on the prevalence of true interactions; **Table S6** (FDR_by_prevalence) therefore tabulates implied FDR across assumed prevalences. At the deployed iLIS threshold (10% FPR; iLIS ≥ 0.223), the implied FDR falls from 93% when true interactions make up 1% of the scored set, to 55% at 10% and 35% at 20%; tightening to 1% FPR (iLIS ≥ 0.551) reduces the corresponding FDR to 72%, 19%, and 10%, with sensitivity dropping from 73% to 38%.

To test the deployed cutoffs without recalibration, each metric’s fixed 10% FPR cutoff was applied to the held-out controls (4,620 pairs: ELM proteins paired with GFP, Wg, or shuffled partners, and shuffled PRS pairs) and held-out positives (330 ELM interactions), and the achieved FPR and sensitivity were reported with Wilson 95% confidence intervals by control class, species, and pLDDT subgroup (**Figure S4B**; **Table S6**). The same fixed AF rank-1 cutoffs were applied without recalibration to the dedicated receptor ECD negative controls (2,105 pairs across all 421 receptor ECD isoforms; see Candidate pair selection), reporting achieved FPR with Wilson 95% confidence intervals across all seven metrics by partner and complex pLDDT subgroup (**Figure S16**; **Table S6**; **Table S4**, sheets Q2 and Q3).

Paired comparisons between metrics at matched FPR used stratified bootstrap resampling (2,000 replicates, one shared set of resampling indices for all metrics, cutoff re-derived within each replicate as the score of the k-th highest control); AUC differences were tested by the paired DeLong method (**Figure S4A**; **Table S4**, sheets F and F2). Sensitivities at matched FPR by interface engagement (**Figure 1D**, **right**; **Figure S3I-J**) were read from the ROC curve by linear interpolation at the target FPR. Discordant calls between iLIS and each other metric were summarized by the likelihood ratio (fraction of positives in the discordant group divided by the fraction of controls), equivalent to precision at equal prevalence, with bootstrap confidence intervals.

Thresholds were computed at 0.1%, 0.5%, 1%, 5%, and 10% FPR across pLDDT subgroups (0 to 50, 50 to 70, 70 to 100, stratifying by how much of each full-length pair is predicted with low confidence) and the total set, for both best-model (AF rank-1) and average-model (five-model mean) scoring (**Table S6**; the pair-level values used for calibration are listed in the Calibration_subset sheet of **Table S1**). Here, pLDDT refers to the model-level confidence reported by AlphaFold-Multimer for the predicted complex as a whole, not a per-chain average. With 1,255 calibration controls, the nominal 0.1% FPR tier is first reached at 2 false positives (achieved FPR 0.16%, used for its tabulated FDR in **Table S6**). For downstream classification, we adopted the best-model total iLIS threshold at 10% FPR (≥ 0.223); the database and network viewer expose the 10%, 5%, and 1% FPR tiers. These thresholds may require adjustment for specific biological contexts; for example, weaker or more transient interactions may benefit from a more permissive threshold, while high-confidence applications may warrant stricter cutoffs.

### Evaluation against *Drosophila* experimental PPI datasets

Predictions were evaluated against four experimental datasets: FlyBi Y2H interactions (Tang et al., 2023), DPiM AP-MS interactions (Guruharsha et al., 2011), DPiM2 AP-MS interactions (Bhat et al., 2024), and literature-curated PPIs from FlyBase (Öztürk-Çolak et al., 2024), BioGRID (Oughtred et al., 2021), and MIST (Hu et al., 2018). Recovery rate was defined as the fraction of experimentally reported interactions scoring above the iLIS threshold. The recovery rates and the comparison with ipTM in **Figure 1E-L** use, for each metric, the highest value among the five models of a pair. For background comparisons of dataset recovery, atlas pairs among the same proteins that a dataset did not report were treated as presumed negatives and summarized by Mantel-Haenszel odds ratios stratified by the higher-degree protein of each pair (degree being the number of atlas pairs that include the protein); because these pairs were nominated by other campaigns and may include true interactions, their positive rate is an upper bound on the background, which makes enrichment odds ratios conservative but could exaggerate an apparent depletion such as that of DPiM2.

### Comparison with computational PPI prediction datasets

For comparison with Peng and Zhao (2026), we obtained Supporting Information S1 Appendix (DOI: 10.1371/journal.pcbi.1013899.s011), which reports pair identifiers, pDockQ, and iLIS for 27,711 AlphaFold-Multimer predictions on *Drosophila* protein pairs derived from STRING ≥ 0.9 confidence. Pairs were canonicalized to FBgn-FBgn pairs and isoform-level duplicates collapsed by maximum score per pair (27,683 unique pairs). Pairs called positive by iLIS (≥ 0.223) but with pDockQ < 0.23, corresponding to the “possible” tier defined by Peng and Zhao (pDockQ < 0.23 with ipTM ≥ 50 on their 0 to 100 scale, equivalent to 0.5, or minimum inter-chain PAE ≤ 10 Å), were tabulated as PPIs uniquely captured by iLIS and each was checked against the minimum inter-chain PAE ≤ 10 Å criterion.

For comparison with the AlphaFold Database heterodimer resource (Han et al., 2026), all 138,938 *Drosophila melanogaster* (taxon 7227) heterodimer models in the bulk archive (138,053 entries in the release summary) were scored directly with the iLIS pipeline (lis.py) and mapped from UniProt accessions to unordered FBgn pairs (137,969 unique gene pairs, of which 2,054 pass the AlphaFold Database high-confidence threshold of ipSAE ≥ 0.6 and pDockQ2 ≥ 0.23, recorded in the passes_quality_threshold flag of the bulk metadata). Across the 29,685 shared gene pairs with model-level outputs in FlyPredictome, agreement was evaluated both under matched single-model selection (AF-Multimer model_1 in both resources) and against FlyPredictome’s deployed best-of-five iLIS (**Figure S9D-F**; **Table S4**, sheets Y1 to Y3). Restricting the comparison to pairs with identical sequences in both resources gave nearly the same correlation.

For analysis of the Schmid et al. (2025) human predictome, iLIS was computed on the complete set of 1,614,047 AlphaFold-Multimer predictions (KIRC-nominated). Positive calls were defined at iLIS_best ≥ 0.223 (10% FPR). SPOC scores for the same pairs were obtained from the predictomes.org S3 deposit (https://predictomes-hsbps-dataset.s3.us-east-1.amazonaws.com/20251110_hs_predictome_pair_scores.csv.gz). To assess concordance with the SPOC classifier (Schmid and Walter, 2025), we compared iLIS positive calls to SPOC ≥ 0.86, the threshold reported by Schmid et al. (2025) for high-confidence calls (10% FDR; **Table S7**).

For the cross-species conservation analysis (**Figure S9H-J**), FlyPredictome positive heterodimers (n = 144,779) and Schmid et al. (2025) positive human heterodimers (n = 225,749; iLIS_best ≥ 0.223 or SPOC ≥ 0.86, resolved to gene symbols) were mapped between species using DIOPT (Hu et al., 2011) fly-human orthology, taking all reported orthologs (many-to-many) and applying the same rule in both directions. A positive pair in one resource was scored as positive in both when any orthologous pair was positive in the other resource, modeled-but-negative when an orthologous pair was scored but below threshold, not-modeled when no orthologous pair was present in the other resource’s scored set, and no-ortholog when either protein lacked an ortholog. Because orthology is many-to-many, the fly → human and human → fly overlaps differ in absolute number. For the ortholog-confidence panel (**Figure S9H**), each fly positive with a modeled human ortholog was binned by the minimum over its two proteins of that protein’s maximum DIOPT score (the number of independent tools supporting the ortholog assignment); within each bin, the fraction also positive in the human predictome (positive for any orthologous human pair, as above) was computed.

### Experimental-evidence stratification

FlyBase physical-interaction records were obtained from the MITAB release fb_2024_02 and mapped to FBgn pairs by PSI-MI method annotation. Because FlyBase curates only a subset of the large high-throughput screens, the three proteome-scale screens were instead taken from their complete reported sets (FlyBi high-throughput Y2H, n = 8,773; DPiM AP-MS, n = 10,103; and DPiM2 AP-MS, n = 21,166), while all remaining low-throughput methods used the FlyBase-curated pairs. iLIS values for every pair were drawn from the full prediction set. Positive rate was computed as the fraction of pairs with iLIS ≥ 0.223, stratified by experimental method, by the number of supporting experimental records, and by the number of distinct supporting methods.

### Missense allele enrichment analysis

Missense mutations were compiled from two FlyBase sources (Öztürk-Çolak et al., 2024; release fb_2025_05): allele descriptions and genetic interaction records. Mutations were mapped to AFM-predicted structural models by exact position matching or, when isoform numbering differed, by alignment-based isoform correction that transfers the mutated position through the longest matching blocks between the FlyBase isoform and the modeled sequence (difflib.SequenceMatcher). Because isoform differences and annotation discrepancies can shift residue numbering, only mutations whose reference amino acid matched the AFM model at the mapped position were retained. Synonymous substitutions (identical reference and alternative amino acids; n = 10) and non-*Drosophila* profiles were excluded, leaving 2,504 unique mutated gene-position pairs across 861 proteins (688,568 residues).

For each residue in a protein, two interaction frequency metrics were computed across all partner models that contain a confident interaction domain (at least one LIR residue): the LIR percentage (fraction of partner models in which the residue falls within the confident interaction domain) and the cLIR percentage (fraction in which the residue makes direct intermolecular contact). Residues were binned into five groups: Zero (never in any interaction domain), and quartiles Q1-Q4 of the non-zero LIR distribution. In 36 very long proteins, the stored LIR residue-list string exceeded the database character limit and was truncated while the shorter cLIR string was preserved; the 2,916 residues with cLIR > 0 and recorded LIR = 0 were retained as contact-positive in category and regression analyses and excluded from within-LIR > 0 subsets, and a further 2,672 residues of these proteins have an undercounted LIR frequency (5,588 of 688,568 residues, 11 of 2,504 mutations). Five single-partner proteins (18 mutations) were retained conservatively (excluding them yields 7.0× versus 6.9× LIR quartile-4 over zero enrichment).

The Cochran-Armitage trend test was used to assess monotonic enrichment of mutation rate across LIR/cLIR bins. Within-gene permutation tests (100,000 permutations) shuffled mutation positions within each gene independently to control for gene-level ascertainment bias. Evolutionary conservation was assessed using PhyloP scores (Pollard et al., 2010) from the 27-way insect species alignment (23 *Drosophila* species and 4 outgroup insects) available through the UCSC Genome Browser (Kent et al., 2002). Logistic regression was performed with mutation status as the dependent variable and cLIR and PhyloP as independent variables. To control for residue burial, relative solvent accessibility (RSA) was computed for each residue from its monomer model in the AlphaFold Protein Structure Database (*Drosophila melanogaster* proteome), with model residues matched to the profile sequence by pairwise alignment rather than by residue number, as the residue’s solvent-accessible surface area (Shrake-Rupley algorithm) normalized to the residue-specific maximum of Tien et al. (2013), and the enrichment was recomputed both restricting to solvent-exposed residues and with RSA included as an additional covariate in the logistic regression (**Figure S11E-G**).

### Prediction-confidence and MSA-depth covariates

For the 852 proteins with a matching AlphaFold Protein Structure Database monomer model (623,354 residues, 2,426 mapped mutations), mutation status was modeled by logistic regression on standardized cLIR or LIR frequency (or their binary forms), adding residue pLDDT of the monomer model, relative solvent accessibility, PhyloP, and LIR frequency singly and jointly, with or without one intercept per gene (gene fixed effects, fitted by Newton-Raphson; genes without any mutation carry no information and drop out). The residue-frequency profiles do not store per-residue inter-chain PAE, so LIR frequency, which is defined by the PAE ≤ 12 Å window, served as the local-confidence covariate.

Because original prediction-time MSAs were not retained, per-column alignment depth (non-gap homologs) and Neff (80% identity reweighting) were recomputed uniformly across all 861 proteins from MSAs regenerated with colabfold_search (ColabFold 1.5.5, MMseqs2 15) against the same databases (UniRef30 2202 and ColabFold environmental 202108) and search settings. To obtain pLDDT and solvent accessibility from the same alignments, each protein was refolded as a monomer from its regenerated MSA (ColabFold 1.6.3, AlphaFold2 monomer weights [alphafold2_ptm], five models, three recycles, no templates, models ranked by pLDDT), and the top-ranked model was mapped to profile positions by sequence identity. Folding completed for 859 of 861 proteins; the two longest (ft, 5,147 residues; hiw, 5,233 residues) were not completed. The completed models covered 678,131 residues (678,124 with PhyloP coverage) and 2,494 mutations, versus 623,361 residues (623,354 with PhyloP coverage) in the database models (absent for 9 proteins and < 90% coverage for 50). Odds ratios are per standard deviation with Wald 95% confidence intervals (**Figure S12**; **Table S4**, sheets W to W5).

### Phenotype severity and phosphosite enrichment analyses

Phenotype severity was assigned from FlyBase genotype-phenotype records (release fb_2025_05) for the 1,998 exact-mapped alleles with linkable annotations; excluding “other” (n = 202) and “unclassified” (n = 476) entries (**Figure S11K**) left 1,320 severity-graded alleles (lethal, semi-lethal, viable/mild). Enrichment of each class at direct-contact residues was tested by one-sided binomial test against the expectation from the same proteins (the mean, over the alleles of the class, of the fraction of contact residues in each allele’s protein); the pooled residue background (64% in domain, 30% at contact) is shown in the figure for reference. Curated phosphorylation sites were obtained from UniProt/Swiss-Prot (reviewed entries, retrieved July 2026) and mapped to the same models; phosphosites were compared with the other Ser/Thr/Tyr residues, carrying no curated phosphosite, of proteins with a curated modification site, and enrichment at direct-contact residues was assessed by two-sided Fisher’s exact test.

### Candidate pair selection

FlyPredictome was assembled from targeted prediction campaigns rather than from an all-by-all screen. Each campaign defined one or two protein pools and a pairing rule; per-campaign pools, pairing rules, input sequence types, and counts are given in **Table S8** and **Table S4** (sheet L). (i) Secreted ligand-receptor screen: 2,408 secreted proteins (from the GLAD secretome lists [Hu et al., 2015] and DeepLoc 2.0 extracellular predictions [Thumuluri et al., 2022]; full-length longest isoforms, signal peptide retained) were each paired with the extracellular domains (ECDs) of 425 isoforms of 256 single-pass transmembrane receptors (collected from UniProt single-pass transmembrane annotations and FlyBase isoforms, and trimmed to predicted extracellular regions excluding N-terminal signal peptides and transmembrane or intracellular segments using DeepTMHMM; Hallgren et al., 2022), in a cross join, of which about 90% was run (927,368 isoform pairs; about 573,000 unique gene pairs).

(ii) Signaling-enzyme screens: about 255 kinases and about 112 phosphatases were each paired with about 1,000 transcription factors (213,308 and 100,119 pairs), and 263 kinases were paired with each other (32,997 pairs). (iii) Reported interactions: every pair recorded in FlyBase, BioGRID, or MIST, including genetic-interaction-only pairs (about 79,700 pairs), and the DPiM, DPiM2, and FlyBi datasets (about 43,600 pairs). (iv) Pathway and organelle sets: all-by-all pairings within KEGG pathways (Kanehisa et al., 2023) or curated organelle lists (about 23,900 pairs). (v) Kinase-centered and collaborator-nominated candidate lists (about 46,000 pairs). Apart from the receptor ECDs and a few early pilot sets, each protein was modeled as its longest annotated isoform at full length, and pairs already predicted in an earlier campaign were not resubmitted. The numbers above are the sizes of the candidate sets, and a pair nominated by more than one campaign is counted in each; the counts in the Results and in **Table S8** assign every unique pair to a single campaign of record and therefore do not map one-to-one onto these sets.

To evaluate the full-length-calibrated thresholds directly on truncated ECD inputs, the 421 sequence-validated receptor ECD isoforms (excluding 4 of the 425 isoforms whose annotated ECD string did not match the parent isoform sequence) were paired under identical AlphaFold-Multimer v2.3 settings (five models, unpaired_paired MSAs, no templates) with the 291-residue tagged GFP construct of the calibration controls and four cytosolic *Drosophila* metabolic enzymes selected for confident cytoplasmic localization and absence of hub behavior (Gnmt, Gpdh2, Gmer, and CG17028; DeepLoc Cytoplasm score > 0.8, 284 to 358 residues, zero reported MIST interactions with the receptors), yielding 2,105 control pairs (18 to 2,884 ECD residues) (**Figure S16**; **Table S6**; **Table S4**, sheets Q to Q3).

For analysis, every prediction record was mapped to a FlyBase gene identifier (FBgn), pairs were treated as unordered, and self-pairs were analyzed separately as homodimers. Each gene pair was assigned a single iLIS value, the maximum over its five models and over all isoform and campaign records of that pair, reducing 1,506,003 heterodimer prediction records (plus 13,818 homodimer records; 1,481,617 records with single-model outputs in **Figure S7C**) to 1,108,443 unique heterodimeric protein pairs and 13,610 unique homodimers (**Table S8**). Interaction calls apply this highest-iLIS value to thresholds calibrated on the AF rank-1 model of each benchmark pair.

Applied to the calibration set, using the highest-iLIS model instead of the rank-1 model changes the achieved FPR at iLIS ≥ 0.223 from 10.0% (95% CI 8.4 to 11.7%) to 10.4% (8.8 to 12.2%) and sensitivity from 72.8% to 74.5% (and from 3.3% to 3.7% on the dedicated receptor ECD controls); 4.5% of positive heterodimers (6,420 of 143,673 with a rank-1 model) are positive only through a model other than rank 1. Taking the maximum over isoforms adds further multiplicity for proteins modeled as several isoforms (in the ECD screen, the gene-level positive rate on the rank-1 model is 11.6% compared with 9.3% for individual rank-1 isoform pairs).

### Prediction-set composition

Each predicted pair was assigned to the source category from which it originated (previously reported PPIs, kinase interactome, secreted ligand-receptor pairs, subcellular-organelle sets, large-scale proteomics datasets, and combinatorial, pathway, or other targeted sets, the last including the phosphatase × transcription factor screen) by the prediction source file recorded for each pair. Each pair was additionally annotated by evidence class: fly literature (a curated physical or genetic interaction, or a pair drawn from a reported interaction dataset, namely FlyBase, BioGRID, DPiM, DPiM2, or FlyBi), interolog (a conserved interaction in another species), or unreported (neither), with the breakdown by source given in **Table S10**.

Interolog support was assessed by mapping both proteins of a pair to their orthologs in other species with DIOPT (Hu et al., 2011) and testing whether an orthologous pair is a reported interaction in that species (MIST cross-species PPI records; Hu et al., 2018). A pair’s interolog score is the maximum, over all species and orthologous pairs that reproduce the interaction, of the lower of the two orthologs’ DIOPT scores; pairs reaching a score of 2 or more were annotated interolog-supported (the PPI network construction used the stricter cutoff of 4).

Counts and positive rates by selection campaign of record are given in **Table S8**, while **Tables S9 and S10** classify the 144,779 positive heterodimers by single primary source (grouping the phosphatase × transcription factor screen under Kinase/phosphatase interactome). Proteome coverage was computed against all unordered pairs among the 13,968 protein-coding genes annotated in FlyBase (release fb_2025_05).

To evaluate external support in the unreported class (**Table S4**, sheet N), unreported positives (101,190 pairs lacking MIST or DIOPT ≥ 2 support at prediction level; 101,125 in **Table S9**) were compared against sub-threshold unreported pairs (731,560 pairs) for shared specific GO cellular-component terms (≤ 50 annotated fly genes), GO protein-complex terms (GO:0032991 descendants), STRING v12.0 coexpression and text-mining associations (scores ≥ 150; Szklarczyk et al., 2023), and recent BioGRID 5.0.262 or FlyBase FB2026_03 physical-interaction records, reporting rate ratios with Wald 95% confidence intervals overall, by FPR tier, and with Mantel-Haenszel stratification by higher-degree protein.

### PPI network construction and clustering

The evidence-supported PPI network was constructed by selecting positive predictions (iLIS ≥ 0.223) supported by either fly literature evidence or conserved interactions in other species (DIOPT ortholog score ≥ 4; Hu et al., 2011). Proteins with fewer than two connections were removed, and the giant connected component was extracted.

Community detection was performed using the Leiden algorithm (Traag et al., 2019; leidenalg Python package, RBConfigurationVertexPartition, resolution = 1.1, seed = 42). Sub-clustering was performed on each main cluster independently using the same algorithm with per-cluster resolution optimization: for each cluster, resolutions from 0.5 to 4.0 (step 0.1) were tested, and the resolution yielding the highest modularity was selected. The significance of each split was assessed by a permutation test: node-to-community assignments were shuffled 100 times while preserving community sizes, and the split was accepted only if the observed modularity exceeded the 95th percentile of the null distribution (P < 0.05). Recursive nested sub-clustering applied the same criterion with a minimum of five proteins.

Gene Ontology enrichment analysis was performed using the hypergeometric test with FlyBase GO annotations (GAF release 2026-03-28), querying Biological Process (GO:BP), Molecular Function (GO:MF), and Cellular Component (GO:CC) at a significance threshold of 0.05 (Benjamini-Hochberg [BH] correction). Redundant terms were removed by Lin semantic similarity (cutoff 0.7). Full cluster assignments, GO enrichment, network edges, and evidence composition are provided in **Tables S11-S15**. An interactive network viewer is available at https://flyark.github.io/FlyPredictome-network.

### Higher-order complex modeling

Selected sub-clusters from the PPI network were folded as complete assemblies using AlphaFold3 (Abramson et al., 2024) via the AlphaFold Server, submitting the sub-cluster member proteins (or, for large sub-clusters, the subunit set resolved in a reference cryo-EM structure) as a single job. The members of a sub-cluster are proteins connected by evidence-supported iLIS-positive predictions; some assemblies also include subunits from outside the module or the network (**Table S16**). For most sub-clusters, we attempted several assemblies, including membership variants generated by adding or removing one or a few subunits at a time (for example, removing subunits that made no contacts in an initial fold and refolding the remaining set; median six attempts per sub-cluster), and retained those that folded into a single connected complex.

Of 3,793 AlphaFold3 jobs whose subunits all belonged to one network cluster, 2,859 returned models, and 970 folded into a single connected complex on the top-ranked model (subunits connected by iLIS ≥ 0.223 with cLIA ≥ 5, or by contact interaction area cLIA ≥ 50, and clash fraction < 0.15); the catalog holds 398 non-redundant single-cluster assemblies (**Table S16**): 385 from these connected jobs after removal of subsets and duplicates, and 13 that are connected for most but not all subunits and are flagged as partial in **Table S16**. Most of the 398 published assemblies are subsets or variants of their sub-cluster rather than the complete module (7 of 398 contain exactly a sub-cluster’s or sub-sub-cluster’s members). Submission order favored candidate subunit combinations with stronger pairwise support, and assemblies were chosen for the catalog by connectivity and non-redundancy, with some added manually, so the catalog is a curated set and not a uniform sample of possible subsets; every folded job is included in the funnel above.

For scoring against experimental structures, we included every assembly whose reference cryo-EM structure covered at least 80% of the network-assigned subunits at the reference-search step (by ortholog mapping; the subunit_coverage column of **Table S16** is recomputed after chain matching and is below 0.8 for 25 of the 202 evaluated assemblies). Because most references are orthologous non-fly structures, predicted and reference chains were matched by normalized BLOSUM62 local alignment (Bio.Align.PairwiseAligner; minimum similarity 0.12) rather than by exact k-mer sharing, which fails on evolutionarily distant subunits. Agreement was quantified without superposition, by subunit-to-subunit contact recovery (a contact defined as any inter-subunit Cβ-Cβ distance ≤ 8 Å, with Cα used for glycine) so that differences in inter-subunit orientation do not affect the measure; recall and precision were computed over the subunits shared between prediction and reference. Because the most divergent complexes are also the least likely to be scorable against any reference, the reported recovery rate is likely an overestimate; the complete selection funnel is given in **Table S4** (sheets O and P; 3,793 jobs to 398 assemblies) and **Figure S25A** (398 assemblies to 178).

For the origin recognition complex, the prediction was additionally superimposed on the experimental structure for display (matchmaker in ChimeraX; Pettersen et al., 2021), and agreement was quantified by whole-complex Cα RMSD and TM-score from multi-chain alignment in US-align (Zhang et al., 2022) and by per-subunit TM-scores from US-align alignment of each subunit to its experimental chain. We further evaluated how each interface-confidence metric scores subunit pairs within these assemblies, comparing at each metric’s 10%, 5%, and 1% best-model FPR thresholds (**Table S6**) the fraction of contacting versus non-contacting subunit pairs that exceed the cutoff (**Figure S24**). The 398 retained assemblies, both cryo-EM-supported and structurally uncharacterized, are cataloged in **Table S16**.

### Clustered interaction fingerprints

For a query protein, the contact profile (cLIR) against each positive predicted partner (iLIS ≥ 0.223) was assembled into a partner × residue matrix, and partners were clustered by shared contact residues using hierarchical clustering (cosine distance, average linkage), with the number of clusters chosen by silhouette score. Per-residue contact frequency, the clustered fingerprint, cluster membership, per-partner metrics, and the mapping of contact residues onto the AlphaFold structure are computed in the browser by the clustered Local Interaction Profiler (cLIP) within LIVIA (Kim and Perrimon, 2026; https://flyark.github.io/LIVIA), reachable directly from each protein in the interactive network.

### FlyPredictome database implementation

The FlyPredictome database (https://www.flyrnai.org/tools/fly_predictome) was implemented as a website built with the Symfony framework and hosted on a traditional LAMP stack, with underlying data stored in a MySQL database. FlyPredictome uses the HTML5 canvas element for drawing heatmaps, Mol* (Sehnal et al., 2021) for protein visualization, Vega-Lite for certain plots, and generates SVG files for Pearson correlation graphs. The backend is written in PHP, while frontend views are rendered with the Twig templating engine. jQuery was used for DOM manipulation, DataTables was integrated for displaying tabular data with search functionality, and Bootstrap was employed for layout, formatting, and styling. Icons throughout the interface are provided by the Font Awesome library. Both the web application and its underlying database are hosted on the O2 high-performance computing cluster at Harvard Medical School, maintained by the Research Computing group.

### Visualization

Data visualization and statistical plots were generated using Python (matplotlib, seaborn, matplotlib-venn) and R. Network visualization used igraph and NetworkX. Sankey diagrams were generated using SankeyMATIC (https://sankeymatic.com). Protein structure visualization was performed using UCSF ChimeraX (Pettersen et al., 2021) and Mol* (Sehnal et al., 2021) via LIVIA (Local Interaction Visualization and Analysis; https://flyark.github.io/LIVIA) (Kim and Perrimon, 2026), a browser-based tool for interface confidence scoring and 3D visualization. Structural alignment of the predicted *Drosophila* Fz-Dsh complex with the human DVL2-FZD4 cryo-EM structure (PDB: 8WM9; Qian et al., 2024) was performed using the matchmaker command in ChimeraX. RMSD values were calculated for the confident interaction region (LIR residues) and for the contact interface residues (cLIR residues) separately.

### Statistics and reproducibility

No statistical method was used to predetermine sample size; sample sizes were determined by available benchmark reference interactions, curated FlyBase missense alleles, predicted candidate protein pairs, and deposited post-cutoff structures. All hypothesis tests were two-sided unless explicitly noted (one-sided binomial tests against the within-protein expectation in **Figure S11K**, paired with a two-sided Cochran-Armitage trend test across severity tiers; the within-gene permutation tests are one-sided). Paired metric comparisons used 2,000 stratified bootstrap replicates with shared resampling indices and paired DeLong tests for ROC-AUC; proportion uncertainties are Wilson 95% confidence intervals, and logistic-regression odds ratios carry Wald 95% confidence intervals. The allele enrichment tests and logistic models treat alleles or residues as independent observations. Exact sample sizes (*n*), test statistics, confidence intervals, and *P*-values are reported in the text, figure legends, and **Tables S3, S4, and S6**. Investigators were not blinded during computational analysis; random seeds were fixed in all resampling analyses.

## ACKNOWLEDGEMENTS

We are grateful to the Research Computing Group at Harvard Medical School for GPU access in the Harvard O2 high-performance computing cluster and the Longwood high-performance compute cluster.

During the preparation of this manuscript, the authors used Anthropic’s Claude and Google’s Gemini to assist in generating custom Python/R scripts for the analysis of prediction results and to improve the readability and language of the text. After using these tools, the authors reviewed and edited the content as needed and take full responsibility for the publication.

This article is subject to HHMI’s Open Access to Publications policy. HHMI lab heads have previously granted a nonexclusive CC BY 4.0 license to the public and a sublicensable license to HHMI in their research articles. Pursuant to those licenses, the author-accepted manuscript of this article can be made freely available under a CC BY 4.0 license immediately upon publication.

## AUTHOR CONTRIBUTIONS

A.K. and N.P. conceived this study. A.K. performed the AlphaFold-Multimer and AlphaFold3 predictions, developed the iLIS framework and allele enrichment analysis, constructed the PPI network, and wrote the original draft. A.K., A.C., A.V., J.R., and Y.H. built the FlyPredictome database. M.H. compiled the ligand-receptor protein sets. A.K., Y.H., and N.P. reviewed and edited the manuscript with input from all authors. N.P. supervised the study.

## DATA AVAILABILITY

The FlyPredictome database, including iLIS values and AlphaFold-Multimer models for all 1.5 million predictions, is available for interactive exploration at the FlyPredictome portal (https://www.flyrnai.org/tools/fly_predictome). The interactive viewer for the evidence-supported PPI network is available at https://flyark.github.io/FlyPredictome-network, where predictions can be filtered at 10%, 5%, and 1% FPR thresholds (**Table S6**) and by evidence type (fly literature, interolog conservation, or both). Download of all-rank prediction details, including pairs scoring below the positive threshold, is available through the FlyPredictome portal. iLIS source code is available at https://github.com/flyark/AFM-LIS.

An index of all supplementary tables is provided in **Table S0**. The benchmark dataset is provided in **Table S1**, the comparison of benchmarked confidence metrics in **Table S2**, and the bootstrap benchmarking results in **Table S3**. The per-campaign candidate generation rules, held-out and time-split benchmark results, metric comparisons, unreported-class assessment, assembly selection funnel, and dataset backgrounds, together with source data for figure panels, are provided in **Table S4**, whose README sheet lists each sheet with the figure or analysis it supports. Per-pair values for the time-split complexes are provided in **Table S5**, and the threshold calibration tables in **Table S6**. iLIS-positive pairs from the Schmid et al. (2025) human predictome, with paired SPOC and KIRC scores, are provided in **Table S7**, and the composition of the prediction set by source in **Table S8**. All positive PPI predictions with confidence scores are provided in **Table S9**, annotated with the prediction set from which each pair originated, its evidence class (fly literature, interolog, or unreported), and whether it was retained in the evidence-supported network; their cross-tabulation by selection source and evidence class is provided in **Table S10**. Network clustering results and GO enrichment analyses are provided in **Tables S11-S15**, and the catalog of AlphaFold3 higher-order assemblies in **Table S16**. Higher-order assembly models are available through the network viewer. Experimental structures used for comparison were obtained from the PDB (accessions given in the text, **Table S5**, and **Table S16**). Source Data are provided with this paper.

## CODE AVAILABILITY

The iLIS analysis pipeline (LIS, cLIS, iLIS, LIA, cLIA, iLIA, iLISA, and the residue-level LIR and cLIR profiles) is available at https://github.com/flyark/AFM-LIS. The code of the interactive network viewer is available at https://github.com/flyark/FlyPredictome-network (viewer at https://flyark.github.io/FlyPredictome-network).

